# The epigenetic eraser LSD1 lies at the apex of a reversible erythroid to myeloid cell fate decision

**DOI:** 10.1101/2021.01.13.426552

**Authors:** Lei Yu, Greggory Myers, Chia-Jui Ku, Emily Schneider, Yu Wang, Sharon A. Singh, Natee Jearawiriyapaisarn, Andrew White, Takashi Moriguchi, Rami Khoriaty, Masayuki Yamamoto, M. Geoffrey Rosenfeld, Julien Pedron, John H. Bushweller, Kim-Chew Lim, James Douglas Engel

## Abstract

H3K4Me demethylase KDM1a/LSD1 is a therapeutic target for multiple diseases, including the β-globinopathies (sickle cell disease and β-thalassemia) since its inactivation has been shown to lead to robust induction of the fetal globin genes. Here we examined the consequences of conditional inactivation of *Lsd1* in adult red blood cells using a new *Gata1^creERT2^* BAC transgene. Loss of *Lsd1* activity in mice blocked erythroid differentiation and expanded GMP-like cells, converting hematopoietic differentiation potential from an erythroid to a myeloid fate. The analogous phenotype was also observed in human HSPC, coincident with induction of myeloid transcription factors (*e.g*. PU.1 an*d* CEBPα). Finally, blocking the activity of myeloid transcription factors PU.1 or RUNX1 at the same time as LSD1 reverted myeloid lineage conversion to an erythroid phenotype. The data show that LSD1 promotes erythropoiesis by repressing myeloid cell fate, and that inhibition of myeloid differentiation reverses the lineage switch caused by LSD1 inactivation.

## Introduction

Sickle cell disease (SCD) is a genetic disorder arising from a single nucleotide transversion, resulting in a Glu to Val amino acid change in the human β-globin protein, β^s^. β^s^ subunits comprise equal parts, with a-globin subunits, of the tetrameric human hemoglobin, HbS [α_2_β_2_^s^]. HbS polymerizes in hypoxic red blood cells (RBC) causing RBC sickling, increased fragility and subsequent destruction, thereby leading to pathological organ damage, episodic acute pain and early death in affected individuals(Murayama, 1966; Platt et al., 1994). Clinical studies showed that the pathophysiological severity of SCD negatively correlates with increased levels of fetal hemoglobin (HbF; α_2_γ_2_)(Platt et al., 1994). β-thalassaemia major (TM; aka Cooley’s Anemia, CA) can arise by inheritance of hundreds of different genetic mutations that lead to diminished or altered expression of the adult β-globin gene(Origa, 2017). However, co-inheritance of hereditary persistence of fetal hemoglobin (HPFH) alleles, normally leading to a benign hematopoietic condition characterized by elevated HbF levels in adults, significantly ameliorates SCD and TM disease pathology, making HbF re-activation in adults a focal therapeutic target for treating the β-globinopathies(Dedoussis et al., 1999, 2000; Marcus et al., 1997; Olivieri and Weatherall, 1998; Papadakis et al., 2002; Yu et al., 2020).

The human β-globin locus spans 70 kb on chromosome 11 and is composed of five sequentially transcribed genes arranged in their temporal order of developmental expression(Fritsch et al., 1980). At around the time of birth, the fetal γ-globin genes are gradually silenced and transcription “switches” predominantly to the adult β-globin gene(Stamatoyannopoulos, 2005). Two underlying mechanisms are currently thought to regulate this behavior: the competition model (Choi and Engel, 1988) first suggested that each of the globin genes competes for the *trans*-activating activity of the LCR (Stamatoyannopoulos, 2005). The second rule suggests that globin gene closest to the LCR has the greatest chance to directly interact with the LCR to be actively transcribed, unless the gene that is linearly closer to the LCR is repressed by gene-autonomous silencing(Bungert et al., 1995; Dillon and Grosveld, 1991; Peterson et al., 1996; Raich et al., 1990; Tanimoto et al., 1999; Wijgerde et al., 1995). A more comprehensive understanding for how the two γ-globin genes are mechanistically regulated in erythroid cells would provide crucial insights into how HbF might be reactivated for therapeutic benefit in TM and SCD patients.

Many studies have reported mechanisms that regulate γ-globin autonomous repression/gene silencing in red blood cells. Notably, mutations in the binding sites for multiple transcription factors (TFs) in the vicinity of the γ-globin genes result in a physiologically benign disorder that leads to abnormally abundant HbF in adult red cells called hereditary persistence of fetal hemoglobin (HPFH)(Yu et al., 2020). These TFs include Bcl11a, LRF, GATA1 and an orphan nuclear receptor heterodimer composed of TR2 and TR4(Chen et al., 2008; Martyn et al., 2018; Tanabe et al., 2002; Tanabe et al., 2007). The core co-repressor subunits of both TR2/TR4 and BCL11a complexes include a scaffold protein, NCoR1, which recruits the epigenetic chromatin modifying enzymes DNMT1 and LSD1, among others, to those TF binding sites(Shi et al., 2013; Tanabe et al., 2007; Xu et al., 2013; Yu et al., 2018).

LSD1/KDM1A is a histone modifying enzyme that specifically removes methyl groups from histone H3 lysine 4 and 9 (H3K4 and H3K9)(Hosseini and Minucci, 2017; Magliulo et al., 2018). Inhibition of LSD1 de-represses γ-globin transcription, leading to the generation of >25% HbF in cultures of differentiated human HSPC(Shi et al., 2013), a level of HbF that is widely acknowledged would be of therapeutic benefit to SCD or TM patients. *In vivo* administration of one LSD1 inhibitor, RN-1, to baboons or mice, animal models commonly used to study SCD, robustly re-activated HbF synthesis (Ibanez et al., 2017; Rivers et al., 2016), and significantly ameliorated the pathological morbidities associated with SCD(Cui et al., 2015). Taken together, the data indicate that safe, efficient LSD1 inhibition would be efficacious for HbF induction in humans, and a widely accessible treatment for SCD and TM by millions of affected individuals worldwide.

Given the promising effect of LSD1 inhibitors on HbF induction *in vitro* and *in vivo*, it becomes important to understand in preclinical studies any broader possible effects of LSD1 inhibition by drug candidates. Pan-hematopoietic *Lsd1* deletion using Mx1Cre(Kerenyi et al., 2013) or inducible RNAi-based LSD1 mRNA knockdown(Sprussel et al., 2012) in adult mice impaired erythropoiesis coupled to an expansion of HSPC, demonstrating that LSD1 plays roles in multiple hematopoietic lineages. Therefore, a detailed mechanistic understanding of how LSD1 inhibition specifically affects adult hematopoiesis is critical to address the potential therapeutic benefit of any candidate LSD1 inhibitor.

In this study, we generated an erythroid-specific inducible Cre mouse line by modifying a *Gata1* BAC that would allow us to conditionally inactivate *Lsd1* exclusively in adult murine erythroid cells. During the analysis of erythroid *Lsd1* conditional loss-of-function (CKO) mice, we discovered that LSD1 also plays a critical role in repressing myeloid gene expression: erythroid-specific *Lsd1* inactivation *in vivo* led to a marked expansion of granulocyte-monocyte progenitor (GMP)-like cells in the bone marrow. Furthermore, we discovered that these novel GMPs surprisingly arise from erythroid progenitors, and that PU.1, an important myeloid transcription factor, was induced in colony forming unit-erythroid (CFU-E) cells recovered from *Lsd1*-CKO mouse bone marrow. Similarly, pharmacological inactivation of LSD1 in human CD34+ HSPC induced to undergo erythroid differentiation re-activated myeloid regulatory genes (*Pu. 1, Cebpa* and *Runx1*) and severely impaired erythroid differentiation. Finally, co-inactivation of *Lsd1* and either *Pu.1* or *Runx1* rescued the erythroid differentiation defect associated with LSD1 inhibition. Taken together, the data indicate that LSD1 functional loss impairs erythropoiesis through a reversible mechanism that induces the activation of a myeloid gene expression program in erythroid progenitors, thus leading to an erythroid to myeloid cell fate conversion. Furthermore, the data show that inhibition of the RUNX1/PU.1 myeloid differentiation axis reverts the *Lsd1* LOF-induced block in erythroid differentiation.

## Results

### LSD1 inhibitors induce HbF but block erythroid differentiation

Current LSD1 inhibitors, as exemplified first by tranylcypromine (TCP), are all small molecules that initially showed promising potential for use as SCD therapeutics because of their ability to stimulate fetal γ-globin expression to high levels(Shi et al., 2013). However, all of the commercial inhibitors that we have tested to date exhibit one or more properties that prohibits their use as promising drug candidates. One novel LSD1 inhibitor that we recently synthesized, CCG50, is a small organic molecule that was designed to bind reversibly to the flavin-dependent catalytic center of LSD1 (data not shown), thus differing from the majority of current (irreversible) commercial inhibitors. We showed that CCG50 (referred to interchangeably as LSD1i) efficiently and specifically inhibited LSD1 function with an IC50 of 115 nM (Supplemental Fig. 1A).

To test whether LSD1i administration would, like TCP, also induce abundant γ-globin expression, we tested erythroid differentiation in human CD34+ HSPC using a standard three-phase culture system (Supplemental Fig. 1B)(Giarratana et al., 2005; Giarratana et al., 2011). In this culture system, proerythroblasts first acquire CD71 (transferrin receptor) followed by CD235a (glycophorin A) surface antigens and subsequently lose CD71 expression to finally display only CD235a as mature erythroid cells(Giarratana et al., 2011; Li et al., 2014). A minority of HSPC remains double negative (15%) while most cells had acquired one (14% CD71) or both (70% CD71+ CD235a+) antigens by d7 (Supplemental Fig. 1C). After differentiation induction on d7 (following the removal of hydrocortisone and human IL3), over 90% of control cells treated with DMSO alone expressed both of these erythroid cell surface markers by d11 and d14, while by d18, 20% of the cells had lost CD71 expression to finally become CD235a+ mature erythroid cells (DMSO panels, Fig. 1A).

At the lowest LSD1i concentration tested (120 nM), the flow cytometric pattern of cells at d11 (last column, Fig. 1A) resembled that of d7 controls (Supplemental Fig. 1C) with the majority (65%) of cells remaining double-positive erythroid precursors. From d14 through d18, the majority (95%) of cells remained CD71+ CD235a+ with only 3-4% converting to more mature CD71-CD235a+ cells. In contrast, at the highest concentration of LSD1i tested (1.1 μM), most of the cells (>70%) remained double negative even by d18 (Fig. 1A). Hence, CCG50 exhibits the same concentration-dependent inhibitory effect on erythroid differentiation as also observed with other LSD1 inhibitors (data not shown).

**Figure 1:**
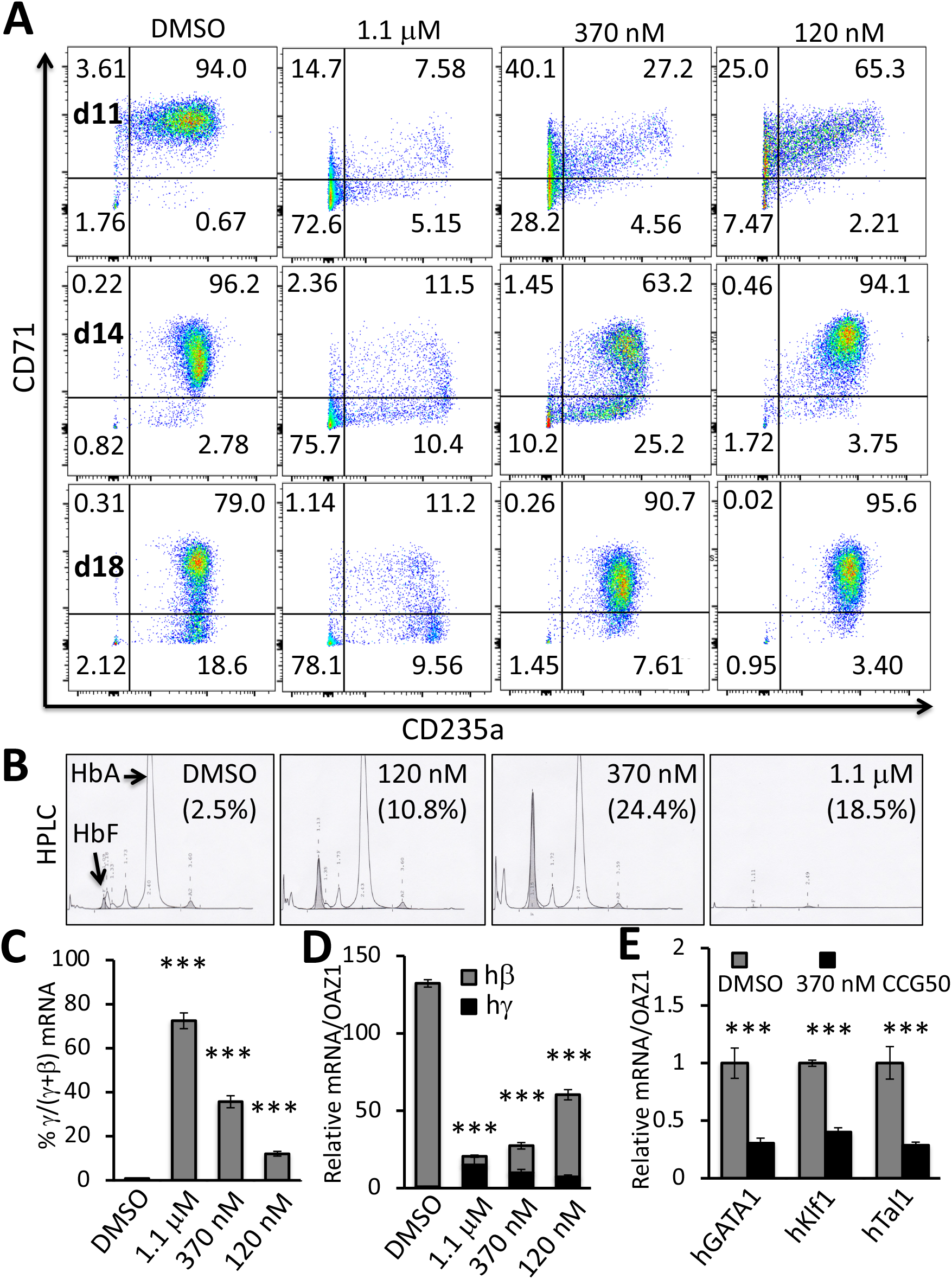
LSD1 inhibition activates γ-globin transcription but blocks erythroid differentiation. (A) Representative flow cytometry plots of human CD34+ HSPC cells undergoing erythroid differentiation after 11, 14 and 18 days (d) in culture. Cells were treated with DMSO or with 1.1μM, 370 nM or 120 nM LSD1 inhibitor CCG050. Cells were monitored for CD71 and CD235a cell surface markers whose acquisition reflect maturing erythroid differentiation stages(Li et al., 2014). Numbers in each quadrant indicates the percentage of gated cells. Results are representative of experiments performed using CD34+ HSPCs from two different healthy adult donors. (B) Representative HPLC chromatograms of d18 cells cultured with or without CCG050 from (A). HbF percentage at each inhibitor concentration is indicated in parentheses. (C) The percentage of γ-globin transcripts in total β-like (γ+β) globin mRNAs at d14. (D) mRNA abundance of γ-globin and β-globin (normalized to OAZ1 internal control mRNA)(Yu et al., 2018) at d14. The γ-globin transcript abundance in DMSO-treated cells was arbitrarily set at 1. (E) Transcript levels of key erythroid transcription factors GATA1, KLF1 and TAL1 (normalized to OAZ1 mRNA) were reduced in d14 cells treated with 370 nM LSD1 inhibitor CCG050. Transcript levels of each mRNA in DMSO-treated cells were arbitrarily set at 1. Data are shown as the means ± SD from three replicates (*** *p*<0.001; unpaired Student’s t-test).

We next assessed HbF synthesis by HPLC analysis of the hemoglobins produced in d18 erythroid cells cultured with or without LSD1i. At all concentrations tested, LSD1i induced γ-globin synthesis and HbF above the baseline level (2.5%) to between 10.8% and 24.4% (Fig. 1B). Globin mRNA analysis indicated that although LSD1i enhanced the γ/(γ+β) ratio in a dose-dependent manner (Fig. 1C), total β-like globin (γ+β) mRNA significantly declined with increasing LSD1i concentration (Fig. 1D). Expression of several key erythroid transcription factors (GATA1, KLF1 and TAL1) was coincidentally reduced after LSD1i treatment (Fig. 1E). Taken together, the data indicate that while low to moderate LSD1 inhibition (at 120 or 370 nM CCG50) robustly induced HbF, higher concentrations impaired erythroid differentiation accompanied by greater cytotoxicity in expanded human CD34+ HSPC.

### Generation of conditional, erythroid lineage-specific deleter transgenic mice

We next investigated whether genetic *Lsd1* CKO would similarly inhibit adult erythroid differentiation in mice. While Mx1-Cre(Kuhn et al., 1995), EpoR-Cre(Heinrich et al., 2004), and Vav1-Cre(Georgiades et al., 2002) deleter strains have been utilized previously to achieve pan-hematopoietic or erythroid-specific recombination of genes *in vivo*, none were ideal for generating conditional LOF alleles (CKO) in adult murine red blood cells, as Cre activity is either not inducible (*e.g*. in EpoR-Cre or Vav1-Cre animals), or is not restricted solely to the erythroid lineage (Vav1-Cre and Mx1-Cre). To circumvent this issue, we generated and characterized erythroid lineage-specific, tamoxifen (Tx)-inducible lines of Cre transgenic mice.

We utilized a 196 kb mouse *Gata1* Bacterial Artificial Chromosome (abbreviated *G1B*) as the delivery vehicle, which we had previously shown to faithfully recapitulate endogenous *Gata1* gene expression(Suzuki et al., 2009; Takai et al., 2013; Yu et al., 2017). A tamoxifen (Tx)-inducible CreER^T2^ fusion gene(Lim et al., 2012), encoding a Cre recombinase whose activity is dependent on the binding of Tx ligand, was inserted at the start codon of *Gata1* by gene editing (Supplemental Fig. 2). Correctly targeted *G1B*CreER^T2^-Neo+ clones were verified by PCR (not shown). The *frt*-flanked Neo selection marker was excised from the BAC, and the resultant modified *G1B*CreER^T2^ BAC clones were verified by PCR and Sanger sequencing (not shown). The BAC was then microinjected into C57Bl/6J zygotes to generate two *G1B*CreER^T2^ transgenic lines, L245 and L259.

To determine the tissue specificity and activity of Cre recombinase in the two *G1B*CreER^T2^ lines, we crossed them to the *Rosa26*-loxP-Stop-loxP-TdTomato (*R26T*) reporter mouse to generate compound *R26T:G1B*CreER^T^ mutant mice, in which TdTomato (TdT) expression is activated by Cre recombinase activity(Madisen et al., 2010). *R26T:G1B*CreER^T2^ mice were injected with Tx on alternate days five consecutive times; untreated *R26T:G1B*CreER^T2^ mice were included as negative controls. Two weeks after the first Tx administration, total bone marrow (BM) cells were collected to analyze TdT epifluorescence in hematopoietic cells. Total BM cells from control and Tx-treated mice were co-stained with anti-CD71 and -Ter119 antibodies to monitor erythroid differentiation.

As erythroid cells mature, they first acquire CD71 expression (gate II; Fig. 2A) before coexpressing both Ter119 and CD71 (gate III) and then gradually lose CD71 immunopositivity (gate IV) to express Ter119 alone (gate V)(Chen et al., 2009). No TdT signal was detected in the BM of either untreated L245 or L259 mice, indicating that CreER^T2^ activity was not “leaky” (blue peaks, Fig. 2B). Since CreER^T2^ was under the transcriptional control of *Gata1 cis*-regulatory elements contained within the 196 kb BAC, we expected that upon Tx administration, only cells with erythroid developmental potential would express TdT. Indeed, fraction I, which primarily represents BM non-erythroid cells, were almost exclusively TdT-negative, whereas > 90% of the cells in fraction II, which are committed to erythroid differentiation, were labeled by TdT (Fig. 2B, red peaks). Notably, the TdT intensity diminished as cells differentiated (fractions III-V), likely due to some combination of global erythroid nuclear condensation, TdT turnover and dilution (Fig. 2B). We also stained whole BM cells with a cocktail of antibodies recognizing non-erythroid hematopoietic cell surface antigens to determine whether CreER^T2^ expression was expressed in B (B220+), T (CD3_ɛ+_) or myeloid (Gr1+, CD11b+) cell lineages. Flow cytometry indicated that very few TdT+ cells were present in the B220/Gr1/CD11b/CD3e+ population (Supplemental Fig. 3, red peak), indicating that the *G1B*CreER^T2^ transgene was active essentially only in the erythroid lineage. Both transgenic lines shared similar TdT expression profiles (Fig. 2B and Supplemental Fig. 3), and therefore only line L259 was used in all subsequent studies.

**Figure 2:**
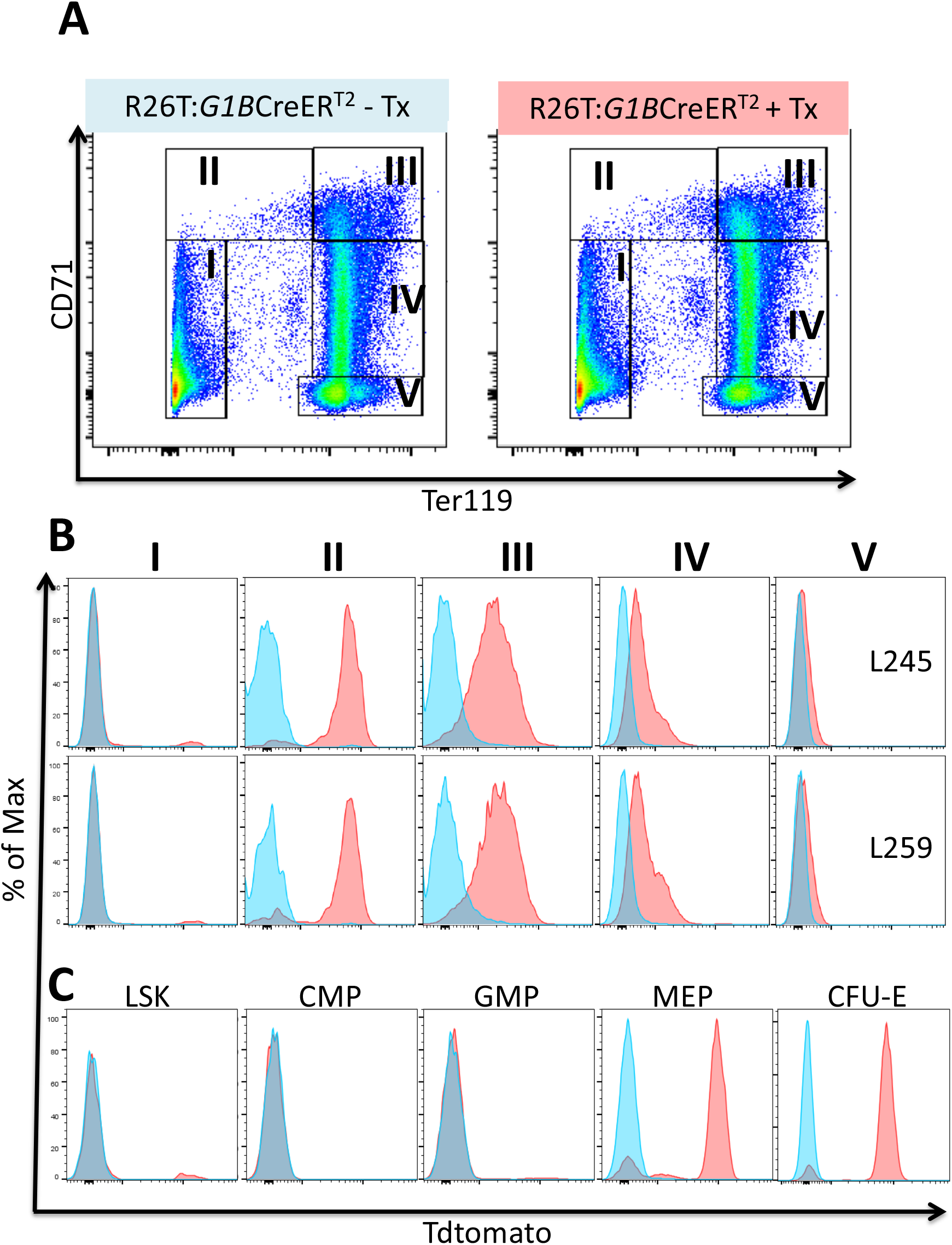
Inducible, erythroid-specific *G1B*CreER^T2^ expression is initiated at the MEP stage. (A) Representative flow cytometry plots showing gating strategies for the anti-CD71 and -Ter119 antibodies used to identify progressively more mature erythroid cells (from stages II to V) among BM cells of untreated (left panel) or 2 mg of Tx (2 mg for 5 times every other day; right panel)-treated *R26T:G1B*CreER^T^ line 259 mice. (B) Representative flow histograms showing TdTomato epifluorescence in cell fractions I-V (as gated in A.) from DMSO-treated (blue peaks) or Tx-treated (red peaks) *R26T:G1B*CreER^T2^ murine transgenic lines L245 (top) or L259 (bottom) bone marrow cells. (C) Representative flow histograms showing TdTomato epifluorescence in the BM Lin-cKit+Sca1+ (LSK), CMP (Lin-cKit+Sca1-CD34+CD16/32-), GMP (Lin-cKit+Sca1-CD34+CD16/32+), MEP (Lin-cKit+Sca1-CD34-CD16/32-) and CFU-E (Lin-cKit+Sca1-CD16/32-CD41-CD150-CD105+) populations (more detailed gating strategies are shown in Supplemental Figs. 5, 6) in untreated (blue peaks) or Tx-treated (red peaks) *R26T*:*G1B*CreER^T2^ (line 259) cells.

Since mouse *Gata1* is expressed in megakaryocytes (Mk), and other myeloid cells(Shimizu et al., 2008), we examined Cre activity in Mk progenitors (MkPs), Mk and mast cells in *R26T:G1B*CreER^T^ mice (Supplemental Figs. 4 and 5). Only 10% of MkP (defined as Lin-cKit+Sca1-CD41+CD150+) (Supplemental Figs. 4A, 4B and 5) and 25% of mature CD41+CD61+ Mks were TdT+ (Supplemental Figs. 4C, 4D). Of the FcRIa+c-Kit+ mast cells, 15% were TdT+ (Supplemental Figs. 4E, 4F). These data suggested that either the *G1*BAC does not contain the regulatory element(s) that are required for *Gata1* expression in Mk and mast cells or that endogenous GATA1 is expressed in only a subset of those lineages. Taken together, the data indicate that the *G1B*CreER^T2^ mouse is a novel, inducible genetic tool that is faithfully expressed in the erythroid lineage.

### Hematopoietic lineage expression of *G1B*CreER^T2^

We next investigated the expression of *G1B*CreER^T2^ within the hematopoietic hierarchy. BM cells were harvested from *R26T:G1B*CreER^T2^ mice that were either injected with Tx or corn oil (control) and then stained with antibodies for Lin-Sca1+c-Kit+ (LSK cells, murine hematopoietic stem and progenitor cells), common myeloid progenitors (CMP: Lin-c-Kit+Sca1-CD34+CD16/32-), granulocyte-monocyte progenitors (GMP: Lin-c-Kit+Sca1-CD34+CD16/32+), megakaryocyte-erythrocyte progenitors (MEP: Lin-c-Kit+Sca1-CD34-CD16/32-) or colony forming unit-erythroid cells (CFU-E: Lin-c-Kit+Sca1-CD41-CD16/32-CD150-CD105+) using conventional cell surface markers to identify each subpopulation(Pronk et al., 2007) (Supplemental Figs. 5–7, Fig. 2C). Of the LSK cells, approximately 5% were TdT+ after Tx-induced CreER^T2^ activation, which may represent an erythroid-or Mk-primed progenitor cell population (Fig. 2C, red peak)(Arinobu et al., 2007). As anticipated, no CMP or GMP cells were labeled after Tx treatment (Fig. 2C). In contrast, 85% of MEP and 90% of CFU-E were TdT+ (Fig. 2C). The data thus indicate that the Cre activity of *G1B*CreER^T2^ mice is detected as early as the MEP stage as well as at all subsequent stages of erythroid differentiation.

### Erythroid progenitor deficiencies in *Lsd1* CKO mice

In previous studies, Kerenyi *et al*.(Kerenyi et al., 2013) reported prenatal lethality as a result of defective erythropoiesis after *Lsd1* deletion. To investigate the effects of *Lsd1* conditional erythroid ablation in adult animals, we examined congenic C57Bl/6 *G1B*CreER^T2^ mice crossed with *Lsd1* homozygous floxed (*Lsd1*^f/f^) mice(Wang et al., 2007). *Lsd1*^f/f^:*G1B*CreER^T2^ (*Lsd1* CKO) or *Lsd1*^+/+^:*G1B*CreER^T2^ (control) mice were administered Tx every other day for two weeks, and mice were sacrificed for hematological analysis 12 hours after the final injection, unless otherwise noted. This strategy maximized the possibility that continuous exposure to Tx would more likely lead to deletion of *Lsd1* in cells that had escaped earlier inactivation.

Peripheral blood analysis indicated that *Lsd1* CKO mice suffered from anemia, as all erythroid parameters were significantly affected (Table 1), while the WBC count was indistinguishable between the *Lsd1* CKO and control groups. Somewhat surprisingly, platelet counts were significantly higher in the *Lsd1* CKO mice (Table 1), consistent with the increase in CD41+ Mks in the BM of *Lsd1* CKO mice (data not shown).

**Table 1:** Differential peripheral blood analysis of *Lsd1*-CKO mice and controls. Mice were IP injected with 2 mg of tamoxifen 7 times on alternate days prior to sacrifice. Data represent the counts of white blood cells (WBC); platelets (PLT); red blood cells (RBC); hemoglobin (Hb); hematocrit (HCT); mean corpuscular volume (MCV); mean corpuscular hemoglobin (MCH). Data are shown as the means ± SD from three to eight mice. (* *p*<0.05; ** *p*<0.01; *** *p*<0.001; unpaired Student’s t-test).

|  | Control | *Lsd1*-CKO |
| --- | --- | --- |
| WBC (x10<sup>3</sup>/μl) | 5.47±1.62 | 5.46±1.41 |
| PLT (x10<sup>3</sup>/μl) | 785±125 | 1032±210* |
| RBC (x10<sup>6</sup>/μl) | 9.76±0.46 | 7.77±0.7** |
| Hb (g/dL) | 12.67±0.4 | 9.66±0.67*** |
| HCT (%) | 43.4±2.71 | 31.12±2.15*** |
| MCV (fL) | 44.47±0.68 | 40.1±1.12*** |
| MCH (pg) | 13±0.26 | 12.42±0.43 |

To investigate the source of the anemia observed in the *Lsd1* CKO mice in greater detail, we performed *in vitro* colony assays of BM cells from *Lsd1* CKO and control mice. The data indicated that colony-forming unit-granulocyte, erythroid, monocyte, megakaryocyte (CFU-GEMM) numbers were indistinguishable between the *Lsd1* CKO and control animals (Fig. 3A). Interestingly, there were slightly, but significantly, more colony-forming unit-granulocyte, monocyte (CFU-GM) colonies in *Lsd1* CKO BM (Fig. 3A). Among committed erythroid progenitor populations, burst-forming unit-erythroid (BFU-E) and CFU-E were quantitatively (by 50% or 90%, respectively; Fig. 3A) and qualitatively reduced (Fig. 3B) compared to control BM, thus indicating a severe depletion in the erythroid progenitor populations as a result of *Lsd1* ablation. Hence, the anemia observed in *Lsd1* CKO mice stemmed from a severe reduction in the number of erythroid progenitor cells.

**Figure 3:**
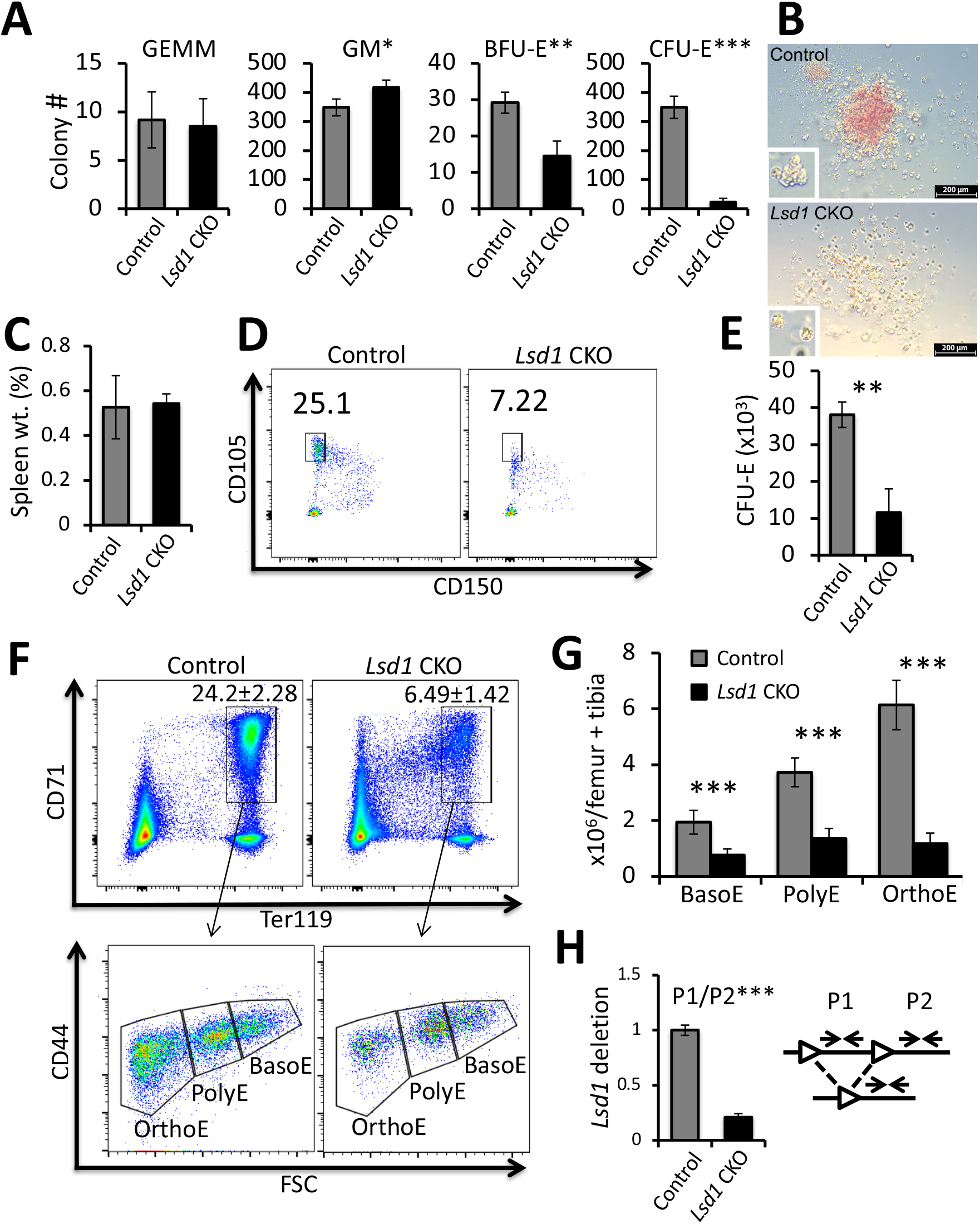
Effects of *Lsd1* deletion in erythroid cells of conditional knockout mice. *Lsd1* CKO and control mice were administered seven Tx IP injections on alternate days. Total BM cells were harvested and processed for colony assays and flow cytometric analyses. (A) Colony numbers of CFU-GEMM, CFU-GM, BFU-E and CFU-E per 10^5^ BM cells of control and *Lsd1* CKO mice. (B) Representative micrographs of BFU-Es from control or *Lsd1* CKO mice taken at low or high (inset) magnification on day 10 of CFU assay. Scale bar = 200 μM. (C) No statistically significant difference is observed in the spleens (as percentages of body weights) of control versus *Lsd1* CKO mice. (D) Representative flow cytometry plots displaying the gating for CFU-Es of control and *Lsd1* CKO mice. (E) Bar graph showing absolute numbers of CFU-E in control and *Lsd1* CKO mice (per two femurs + two tibias). (F) Representative flow cytometric plots showing the gating strategy for defining CD71+Ter119+ erythroid precursor cells (top panels), which were subsequently gated by CD44 staining versus forward scatter (FSC) to separate basophilic erythroblasts (BasoE), polychromatic erythroblasts (PolyE) and orthochromatic erythroblasts (OrthoE)[bottom panels]. (G) Absolute numbers of BasoE, PolyE and OrthoE in control and *Lsd1* CKO BM. (H) Deletion efficiency of *Lsd1* flox alleles in flow-sorted BasoE was determined by qPCR. Primer pair P1 detects *Lsd1* genomic DNA flanked by the two loxP sites while primer pair P2 detects *Lsd1* genomic DNA that is unaffected by Cre-mediated deletion. Data are shown as the means ± SD. (*** *p*<0.001; unpaired Student’s t-test).

Splenomegaly was not observed in *Lsd1* CKO mice, suggesting that erythroid progenitors responsible for extramedullary hematopoiesis were also affected (Fig. 3C). In agreement with the colony assays, flow cytometry showed that the absolute number of CFU-E in *Lsd1* CKO mice was significantly reduced when compared to controls (Fig. 3D, E). Based on AnnexinV staining(Riccardi and Nicoletti, 2006), this reduction was not attributable to increased programmed cell death of CFU-E (Supplemental Fig. 8). The data indicate that *Lsd1* CKO leads to a severe block in generating a normal number of erythroid progenitor cells, most markedly at the CFU-E stage, and consequentially to fewer circulating mature red blood cells, and that this decrease is not due to enhanced erythroid cell death.

In the *Lsd1* CKO BM, the number of CD71+Ter119+ erythroid precursor cells was significantly lower than in control BM (Fig. 3F). CD71+Ter119+ erythroid precursor cells can be further fractionated into progressively more mature erythroblasts (basophilic, BasoE; polychromatic, PolyE; or orthochromatic; OrthoE) using a CD44+ vs. forward scatter (FSC) flow cytometry gating strategy (Fig. 3F)(Chen et al., 2009). The absolute cell number of BasoE, PolyE and OrthoE were all significantly reduced in the *Lsd1* CKO CD71+Ter119+ BM (Figs. 3F, 3G), and the >70% reduction in erythroid precursor cells was likely due to continuous differentiation dysfunction that is first detectable in BFU-E (Fig. 3A, E).

To test the possibility that the residual erythroid precursor cells in the treated animals were not simply cells that had escaped *Lsd1* conditional deletion, we quantified Cre-mediated deletion efficiency in flow-sorted BasoE. The data show that >80% of flow-sorted BasoE had deleted the loxP-flanked DNA (Fig. 3H). Hence, *G1B*CreER^T2^ achieved highly efficient conditional *Lsd1* inactivation in adult erythroid lineage cells.

During erythroid cell maturation, there is normally an exuberant increase in cell numbers as erythroid precursor cells transit from basophilic to polychromatic to orthochromatic erythroblasts (control mice, Fig. 3G, gray bars). However, in *Lsd1* CKO mice, the cell numbers increased only marginally between the basophilic and orthochromatic stages (Fig. 3G, black bars). This difference in differentiation potential between *Lsd1* CKO and control mice indicates that LSD1 plays an important role in maturation at multiple sequential stages during erythroid cell differentiation.

We finally asked whether an increase in programmed cell death might account for the observed deficiency in erythroid precursors in *Lsd1* CKO mice. Early and late apoptotic cells (AnnexinV+DAPI- and AnnexinV+DAPI+, respectively(Yu et al., 2017)) were analyzed in BasoE, PolyE and OrthoE cells in *Lsd1* CKO and control mice. Surprisingly, the late apoptotic frequency of both BasoE and PolyE in *Lsd1* CKO mice was significantly reduced (to approximately half), whereas cell death of OrthoE in *Lsd1* CKO mice was unchanged (Supplemental Fig. 9). Furthermore, the cell cycle in CD71+Ter119+ precursor cells was unaffected (Supplemental Fig. 10). Taken together, the data indicate that adult *Lsd1* CKO leads to a block in the differentiation of erythroid precursor cells in a manner that is independent of programmed cell death or alterations in the cell cycle.

### *Lsd1* CKO erythroid progenitors acquire myeloid characteristics

The data presented thus far indicates that *Lsd1* conditional LOF results in marked depletion of definitive erythroid progenitor and precursor cells that is independent of cell cycle and cell death. Curiously, data from the colony assays showed that the GM colony number significantly increased (Fig. 3A). To potentially explain this curious observation we hypothesized that erythroid progenitor cells might have acquired myeloid features through a lineage switch that reciprocally impaired erythroid potential. To test this hypothesis, we examined the LSK, CMP, MEP and GMP progenitor cell populations in *Lsd1* CKO and control mice (Figs. 4A, 4B and Supplemental Fig. 7); the data show that the absolute numbers of LSK, CMP and MEP were comparable between the groups. However, consistent with the hypothesis that erythroid progenitors were being diverted into other lineages, the GMP population significantly increased in the *Lsd1* CKO BM (Figs. 4A, 4B). Curiously, we also detected prominent expansion of a new Lin-c-Kit+Sca1-CD16/32+CD34-cell population in the *Lsd1* CKO bone marrow (Fig. 4A).

**Figure 4:**
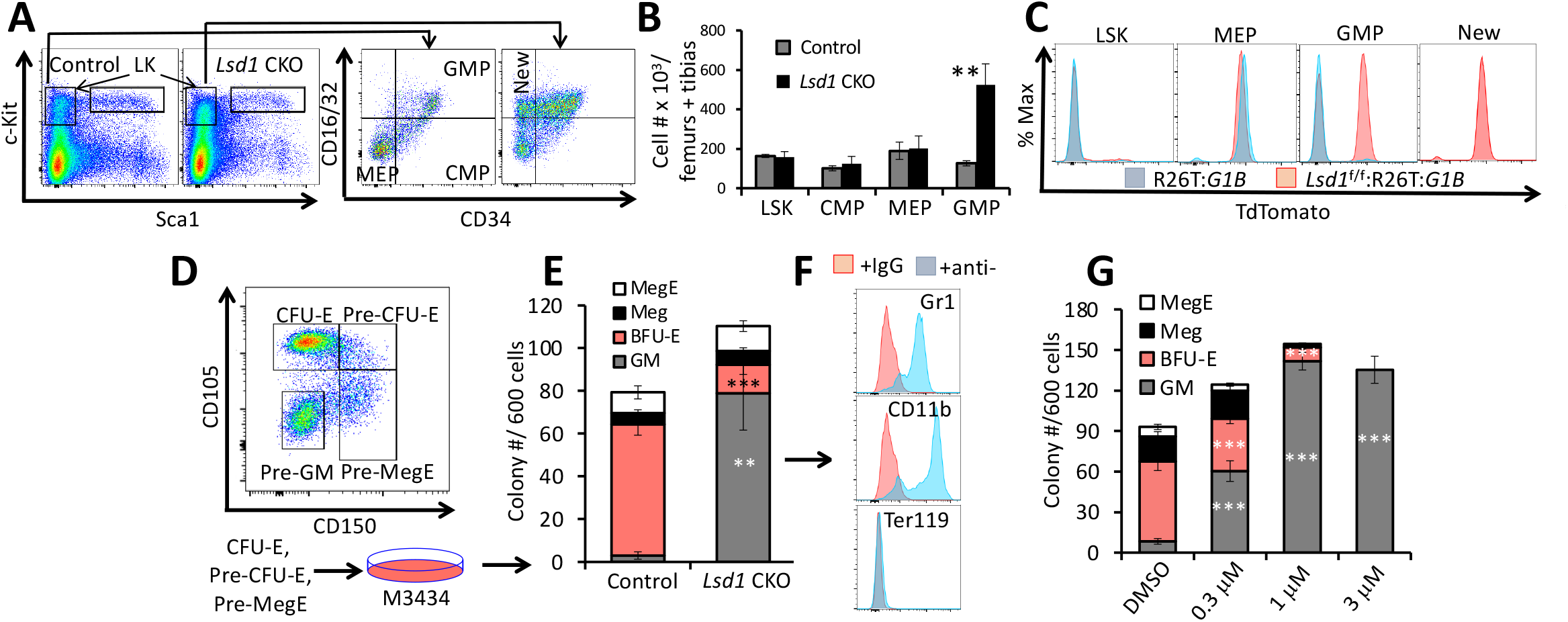
LSD1 LOF shifts erythroid differentiation potential to the GM lineage. Representative flow cytometry plots showing the gating strategies for Lin-Sca1+cKit+ (LSK) cells, CMP, GMP and MEP (A), and their absolute cell numbers (B) in Tx-treated *Lsd1* CKO or control mouse BMs. (C) TdTomato epifluorescence was analyzed in individual cell populations of Tx-treated R26T:*G1B*CreER^T2^ control (blue peaks) or *Lsd1* CKOT (red peaks) mice. (D) Erythroid progenitors give rise to GM colonies in LSD1-CKO BM. Sorted CFU-E, Pre-CFU-E and Pre-MegE from Tx-treated control and *Lsd1* CKO mice were pooled for seeding in colony forming unit assays. Colony types [Meg, mixed MegE, GM and BFU-E (per 600-sorted cells)] were quantified and presented in panel (E) (F) Representative flow cytometry plots of individual GM colonies picked from the *Lsd1*-CKO. The cells were stained with anti-Gr1,-CD11b or-Ter119 antibodies. (G) Colony numbers of Meg, MegE, CFU-GM and BFU-E per 600 sorted cells of control mice treated with DMSO or different concentrations of CCG50 LSD1i. Data are shown as means ± SD from three mice. (** *p*<0.01; *** *p*<0.001; unpaired Student’s t-test).

To test whether the expanded GMP population in *Lsd1* CKO mouse BM might arise from the erythroid lineage origin, we generated *Lsd1*^f/f^:*G1B*CreER^τ2^:*R26T* triple mutant mice (hereafter referred to as *Lsd1* CKOT) in order to lineage trace TdT fluorescence in the expanded GMP population that was revealed in the *Lsd1* CKO mice. Since the erythroid cells, but not GMP, in *G1B*CreER^T2^:*R26T* mice were labeled by TdT (Fig. 2C), we reasoned that the surfeit GMP cell population in *Lsd1* CKOT mice would be TdT+ if they were in fact derived from erythroid precursor cells. As expected, LSK cells in control and *Lsd1* CKOT mice were unlabeled by the lineage marker while MEP cells in both genotypes were TdT+ (Fig. 4C), consistent with earlier observations (Fig. 2C). In contrast, in Tx-treated control mice GMP were not labeled by TdT (Fig. 2C), while 60% of GMP in *Lsd1* CKOT BM were TdT+ (Fig. 4C), which is entirely consistent with the observed three-fold increase in GMP cell number in these animals (Figs. 4A, 4B), Based on detailed characterization of hematopoietic TdT expression (Fig. 2 and Supplemental Figs. 3 and 4), we conclude that these LSD1-deficient TdT+ GMP cells must have originated from an erythroid progenitor population.

We next asked whether the GMP population in *Lsd1* CKO mice (Fig. 4B) might be differentially susceptible to programmed cell death. While the early apoptotic populations of GMP were comparable in *Lsd1* CKO and control groups, late apoptosis was reduced by 50% in the *Lsd1* CKO GMP fraction (Supplemental Fig. 11), suggesting that the ectopic TdT+ GMPs are not differentially susceptible to apoptotic cell death and may simply differentiate into more mature myeloid cells. To test this hypothesis, we flow sorted the CFU-E, Pre-CFU-E and Pre-MegE from control and *Lsd1*-CKO BM and then seeded all three erythroid progenitor populations as a mixture into MethoCult media to test their differentiation potential (Fig. 4D). The majority of these erythroid progenitors in the control mice gave rise to BFU-E, Meg and MegE colonies as well as to a few (possibly contaminating) GM colonies. In contrast, erythroid progenitors from *Lsd1*-CKO mice largely shifted differentiation potential from BFU-E to GM lineages, as the majority of colonies generated from *Lsd1* CKO erythroid progenitors acquired GM characteristics (Fig. 4E). To determine whether the CFU-GM colonies derived from *Lsd1* CKO erythroid progenitors are indeed myeloid cells, we picked individual colonies from *Lsd1* CKO-seeded media and subjected the cells to flow cytometry, examining those cells for the expression of surface markers Gr1, CD11b (myeloid markers) and Ter119 (an erythroid marker). The majority of these individual GM colonies were composed of cells that were Gr1 + or CD11 b+; most significantly, none of the cells expressed the Ter119+ marker (Fig. 4F), indicating that these myeloid colonies arose from early erythroid (CFU-E or Pre-CFU-E) progenitors in Tx-treated *Lsd1* CKO mice.

To test the hypothesis that LSD1 inhibition leads to an erythroid to myeloid cell fate switch (EMS) using an orthogonal strategy, we isolated untreated CFU-E, Pre-CFU-E and Pre-MegE from wild type mice by flow sorting, and then treated them with increasing concentrations of LSD1i in MethoCult. Consistent with the data from genetic *Lsd1* LOF experiments (Fig. 4E), LSD1i treatment of these cultures induced EMS in a dose-dependent manner (Fig. 4G; *e.g*. 3 μM LSD1i treatment fully converts the erythroid to myeloid lineage differentiation potential). Taken together, these data underscore the hypothesis that both *Lsd1* CKO or LSD1 pharmacological inhibition in erythroid progenitor cells leads to an erythroid to myeloid cell fate conversion.

### *Pu.1* is activated at the CFU-E stage in *Lsd1* CKO mice

To better understand the molecular basis for why *Lsd1* LOF in erythroid progenitors leads to acquisition of GMP features, we flow sorted CFU-E cells from *Lsd1* CKO and control mice (Fig. 3D). When we compared gene expression profiles in the two populations, LSD1 mRNA was reduced (as anticipated) by more than 90% in the CKO cells. PU.1 is a well characterized myeloid transcription factor(Laslo et al., 2006; Rosenbauer and Tenen, 2007; Scott et al., 1994), and transcriptional activation of *Pu.1* is regulated by RUNX1 in HSPC(Willcockson et al., 2019). Consistent with the observed increase in GMP activity in *Lsd1* CKO mice, PU.1 expression was 7-fold higher in their CFU-E. In the same cells, GATA1 mRNA was slightly but significantly reduced, another indication of impaired erythroid differentiation, whereas RUNX1 levels were unchanged (Fig. 5A).

**Figure 5:**
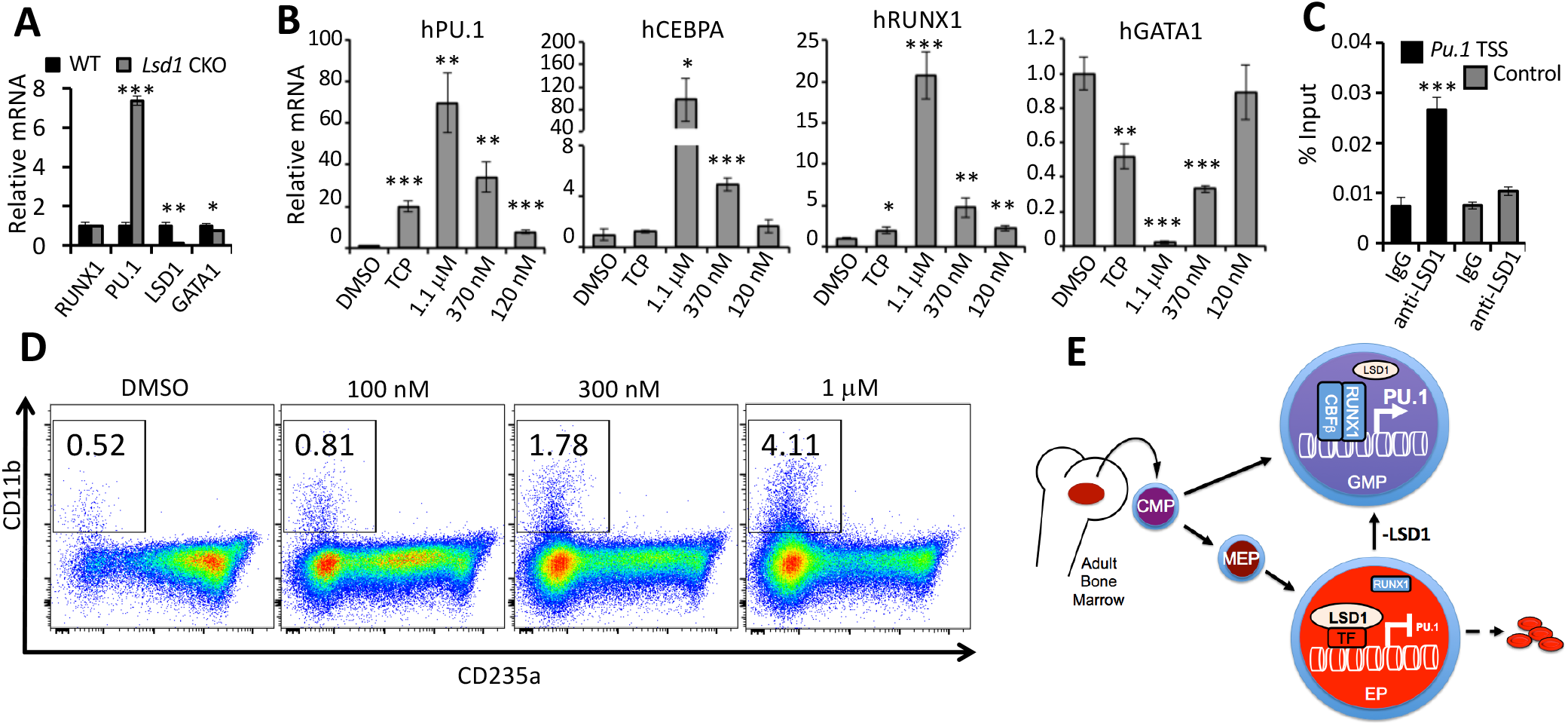
LSD1 directly inhibits myeloid differentiation genes in erythroid cells. (A) Relative mRNA levels of RUNX1, PU.1, LSD1 or GATA1 (normalized to 18s ribosome RNA) in sorted CFU-E cells from Tx-treated control and *Lsd1*-CKO mouse BM. (B) Histograms depicting the relative mRNA levels of human PU.1, CEBPα, RUNX1 and GATA1 (normalized to OAZ1 transcript level) in d14 erythroid cells differentiated from human CD34+ cells at d14 of culture assayed by qRT-PCR. Cells were expanded in the presence of DMSO or LSD1 inhibitors TCP (3 □M) or LSD1i (at three different concentrations). (C) ChIP-qPCR analysis of LSD1 binding to the PU.1 promoter in HUDEP2 cells. Differential enrichment was detected at the PU.1 transcriptional start site (TSS, black bar), but not at an adjacent site included as a negative control (gray bar). Rabbit IgG was included as a negative control. (D) CCG50 treatment enhances CD11b expression during erythroid differentiation induction of human CD34+ cells. Cells were treated with LSD1i (100 nM, 300 nM or 1 μM) or DMSO on day 7. CD11b and CD235a were analyzed at d11 of differentiation. Data are shown as the means ± SD. (* *p*<0.05; ** *p*<0.01; *** *p*<0.001; unpaired Student’s t-test). (E) LSD1 maintains normal erythropoiesis by repressing Pu.1 expression in erythroid progenitors. LSD1 gene deletion or protein inhibition leads to abnormal activation of *Pu.1*, likely through RUNX1, and shifts the differentiation potential from the erythroid to myeloid lineage.

To explore how LSD1 inhibition in erythroid progenitor cells de-represses the expression of common myeloid markers, we performed qRT-PCR on human CD34+ erythroid differentiated cells after 14 days of *in vitro* incubation in the absence or presence of LSD1 inhibitors (Supplemental Fig. 1B). Consistent with the mouse *in vivo* data, two inhibitors significantly activated two myeloid regulatory genes, *Pu. 1* and *Cebpa*, whereas expression of erythroid regulator GATA1 was significantly impaired (Fig. 5B). Notably, treatment with LSD1i significantly induced RUNX1 expression *in vitro* (Fig. 5B) even though *Lsd1* CKO does not affect RUNX1 expression in murine CFU-E *in vivo* (Fig. 5A).

To test whether LSD1 directly regulates *Pu.1* expression, we performed LSD1 ChIP assays in HUDEP2 cells, a human cell line that reflects human adult definitive erythroid cell progenitors(Kurita et al., 2013). The results show that LSD1 is bound at the transcriptional start site of the human *Pu.1* gene (Fig. 5C), an observation consistent with LSD1 ChIP-seq data reported for K562 myeloerythroid lineage cells (<u>UCSC genome browser</u>).

To determine whether LSD1i induces EMS in differentiated human CD34+ cells, we performed erythroid differentiation (adding LSD1i or DMSO treatment beginning at d7) for 14 days (Supplemental Fig. 1B). LSD1i addition enhanced expression of myeloid marker CD11b, up to 4.1% of total, in a dose-dependent manner (Fig. 5D). The efficiency of EMS in the CD34+ erythroid differentiation cultures was less pronounced than in semi-solid media (Fig. 4E-G), perhaps as a consequence of the absence of myeloid cytokines in the erythroid differentiation media.

In summary, these data indicate that LSD1 normally acts to repress *Pu.1* expression in order to maintain unidirectional lineage potential in erythroid progenitor stages. Once *Lsd1* is deleted or LSD1 protein is inactivated by inhibitors, *Pu.1* is induced in erythroid progenitors, likely through increased activity of RUNX1(Willcockson et al., 2019), which initiates the GMP cascade and impairs normal erythroid differentiation (Fig. 5E).

### *Pu.1* or *Runx1* LOF rescues the LSD1i-mediated block of erythroid differentiation

To test whether or not the RUNX1/PU.1 axis plays a functional role in blocking erythroid differentiation that is induced by LSD1 genetic or pharmacological inactivation, we generated *Pu.1* or *Runx1* homozygous knockout HUDEP2 cell clones (Supplemental Figs. 12, 13) to ask whether ablation of either gene could rescue the erythroid differentiation block resulting from LSD1 inhibition. LSD1i treatment at 300 nM CCG50 significantly reduced the fractional percentage of erythroid cells (by 50%) in control HUDEP2 culture (Fig. 6A and Supplemental Figs. 14A, 14B), underscoring the earlier conclusion that LSD1i treatment impairs erythroid differentiation. Remarkably, that reduction cells was rescued by either *Pu.1* or *Runx1* LOF (Fig. 6A and Supplemental Fig. 14). Consistent with the erythroid differentiation block visualized by flow cytometry, LSD1i treatment of control HUDEP2 cells repressed γ-globin, β-globin and GATA1 expression and activated *Pu.1* expression, effects which were rescued in *Pu.1* or *Runx1* knockout HUDEP2 cell clones (Fig. 6B). Taken together, the data indicate that RUNX1/PU.1 activation plays a key role in blocking erythroid lineage differentiation that is induced by LSD1 inhibition.

**Figure 6.**
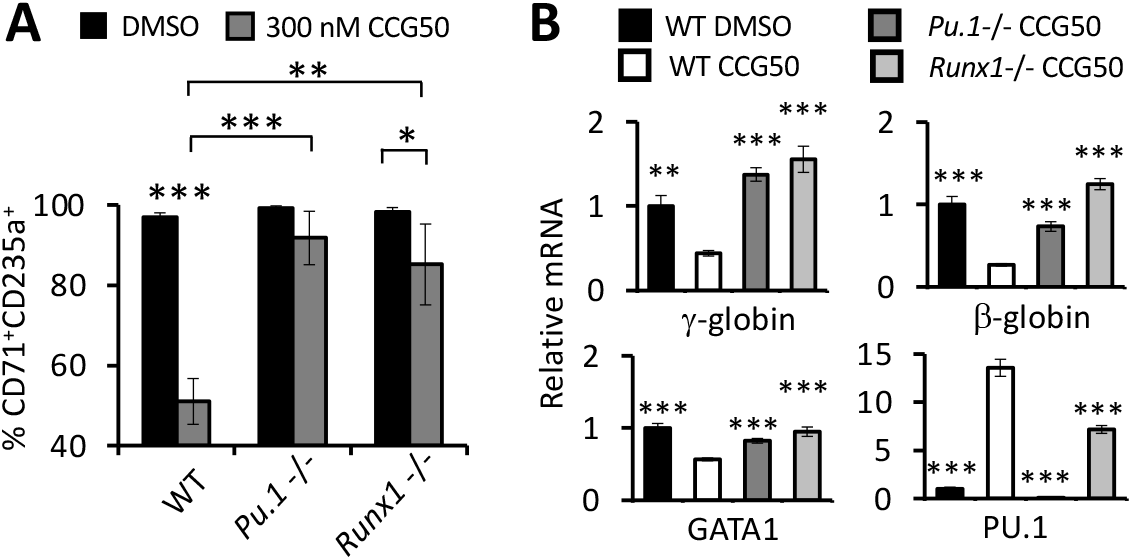
*Pu.1* or *Runxl* genetic knockout rescues the erythroid differentiation inhibition induced by blocking LSD1 activity. **(A)** CD71 and CD235a flow cytometric analysis of three independent non-targeting sgRNA infected cells (WT, left), four *Pu.1*-/-clones (middle) or four *Runx1*-/-clones (right) treated with DMSO (control) or 300 nM CCG50. Data are shown as the average value of percentage of CD71+CD235a+ cells from multiple clones in Supplemental Fig. 14. Impaired erythroid differentiation (as reflected by the reduced percentage of CD71+CD235a+ cells) in the WT group after CCG50 treatment was rescued by either *Pu.1* or *Runx1* knockout. (B) mRNA levels of γ-globin, β-globin, GATA1 or PU.1 in random sgRNA control cells after treatment with DMSO or 300 nM CCG50, or P3-18 (*Pu.1*-/-) and R2-47 (*Runx1-/-*) knockout clones treated with 300 nM CCG50. Data were shown as the means ± SD. (* *p*<0.05; ** *p*<0.01; *** *p*<0.001; unpaired Student’s t-test).

### RUNX1 inhibitors rescue the LSD1i-mediated block of erythroid differentiation

As LSD1 inhibitors have been proposed as potential therapeutic agents for SCD clinical treatment, we next asked if pharmacological inhibition of the RUNX1/PU.1 pathway could rescue the erythroid differentiation blocking effects of LSD1i treatment. As no effective PU.1 inhibitor is commercially available, we tested Ro5-3335, a published RUNX1 inhibitor (RUNX1i) (Cunningham et al., 2012). To investigate whether co-treatment with another, better characterized RUNX1i plus LSD1i could rescue the erythroid differentiation defect caused by LSD1i treatment alone, we also investigated the activity of RUNX1i AI-10-104(Illendula et al., 2016) to the CD34+ erythroid differentiation cultures.

Either RUNX1i was added to CD34+ erythroid differentiation media at the same time as LSD1i, and erythroid cell maturation was examined at d11 by flow cytometry (Supplemental Fig. 1C). Treatment with LSD1i alone inhibited erythroid differentiation as indicated by the reduced production of CD71+CD235a+ cells (Fig. 7A). However, co-treatment of cultures with LSD1i plus either RUNX1i rescued the erythroid differentiation block in a dosedependent manner (Fig. 7A). Although RUNX1i co-treatment rescued the erythroid differentiation block induced by LSD1i, significant cell death was also observed at d11 after Ro5-3335 addition (data not shown). To exclude the effect of RUNX1i-induced cell death during erythroid differentiation, we also analyzed cells at d9, just two days after initiating treatment with the inhibitors (Supplemental Fig. 15A). We found that co-treatment with either RUNX1 inhibitor partially rescued the LSD1i-mediated erythroid differentiation block (Fig. S15B).

**Figure 7.**
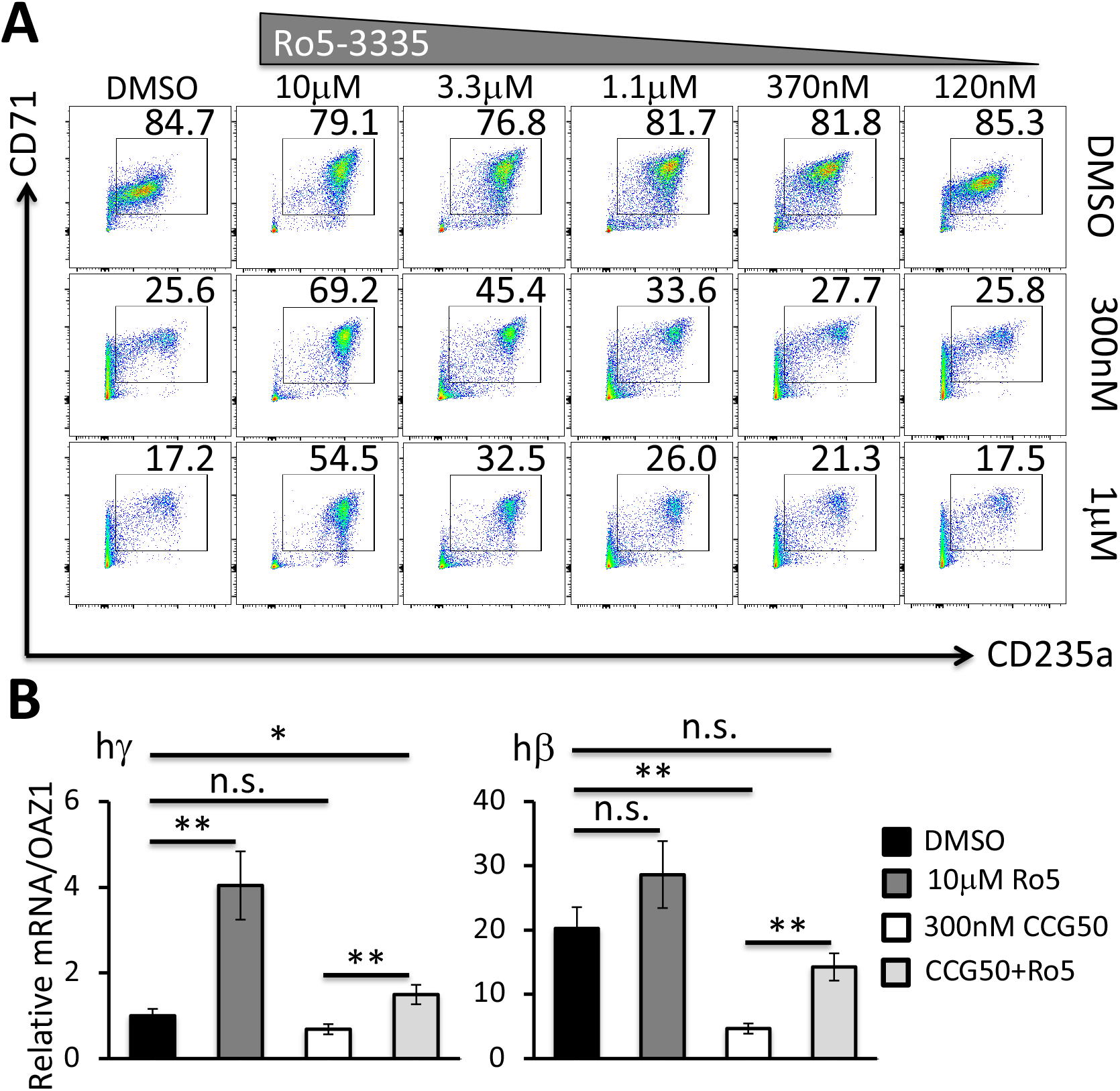
RUNX1 inhibitor co-treatment partially rescues the erythroid to myeloid conversion by LSD1 inhibitors. **Sanger sequencing data Sanger sequencing data (A)** CD71/CD235a staining of human CD34+ cells showing erythroid differentiation at d11, after 4 days of treatment with CCG50 alone (300 nM or 1 μM), RUNX1 inhibitor Ro5-3335 alone (10 μM to 120 nM) or both. **(B)** mRNA abundance of γ-globin and β-globin at d11 after inhibitor treatments. γ-globin transcripts in DMSO were arbitrarily set at 1. Data is shown as the means ± SD. (* *p*<0.05; ** *p*<0.01; unpaired Student’s t-test).

Finally, we examined γ- and β-globin expression after treating the CD34+ erythroid differentiation cultures with RUNX1i, LSD1i or both, at d11 of erythroid-induced differentiation (Fig. 7B). RUNX1i co-treatment with LSD1i significantly rescued globin gene expression that was repressed by LSD1i treatment alone. Surprisingly, γ-globin expression was induced 4-fold by RUNX1i treatment alone, suggesting that RUNX1 may play a previously uncharacterized but LSD1-independent role in fetal globin gene repression in adult erythroid cells (Fig. 7B). Taken together, the data show that pharmacological inhibition of the RUNX1/PU.1 axis partially rescues the differentiation block induced by LSD1i treatment of erythroid precursor cells.

## Discussion

In this study, we initially set out to investigate the effects of LSD1 inhibition on erythropoiesis *in vivo*; this required the generation of a new erythroid lineage-specific, inducible Cre mouse line to explore the effects of conditional *Lsd1* LOF in murine adult erythroid cells. Previously, Takai *et al*. showed that a mouse *Gata1*-GFP knock-in BAC exhibited GFP expression in MEP, that this fluorescence persisted as erythropoiesis progressed, and then gradually weakened in basophilic and polychromatic erythroblasts(Takai et al., 2013). In the *R26T:G1B*CreER^T2^ mice, we observed about 5% of Tx-treated LSK cells were labeled by TdTomato (Fig. X). However, none of the downstream myeloid progeny (such as GMP) were fluorescent, indicating that the GATA1+ LSK cells in these mice do not differentiate into myeloid lineage cells.

The *in vivo* role of LSD1 within the murine hematopoietic compartment has been explored previously. Using the pan-hematopoietic Mx1Cre transgene to inactivate *Lsd1*, Kerenyi *et al*. reported that LSD1 deficiency impacted HSC self renewal(Kerenyi et al., 2013). Furthermore, they showed that *EpoR*Cre-mediated *Lsd1* deletion *in vivo* resulted in prenatal murine lethality due to anemia. Using the *G1B*CreER^T2^ line first reported here, we were able to examine the consequences of *Lsd1* ablation in adult erythropoiesis. We showed that definitive erythropoiesis was compromised as a consequence of drastically reduced erythroid progenitor and precursor cell numbers.

GATA1 was previously reported to transcriptionally repress PU.1 expression by directly binding to GATA consensus sites at the TSS of *Pu. 1* in erythroblasts(Chou et al., 2009). We confirmed LSD1 binding at these same sites in human erythroid HUDEP2 cells by ChIP assays (Fig. 5C). Interestingly, PU.1 forced overexpression impairs erythroid differentiation and leads to erythroleukemia(Moreau-Gachelin et al., 1988). Mice that express GATA1 at 5% of wild type levels also display a propensity for erythroleukemia(Shimizu et al., 2008). Curiously, we detected the expansion of a GMP-like population as well as a new immunophenotype population of Lin-cKit+Sca1-CD34-CD16/32+ cells in *Lsd1* CKO mice (Fig. 4A). Whether these unusual cells might represent harbingers of a pre-leukemic state or a previously undiscovered myeloid progenitor state will required further investigation. In this regard, long term monitoring with an increased propensity for leukemogenesis in the *Lsd1* CKO mice will certainly be of interest.

Increased levels of LSD1 have been detected in different neoplasms, and LSD1 was shown to play a key role in carcinogenesis, highlighting the potential therapeutic effect of LSD1 inhibitors in cancer and myelodysplastic syndrome (MDS) treatment(Fang et al., 2019). However, current LSD1 inhibitors exert multiple side effects, including anemia, thus potentially lessening their utility for targeted therapeutic development(Wass et al., 2020).

In this regard, the mechanisms of LSD1 action revealed in this study may illuminate a path to the development of safe and effective therapeutics for the treatment of the β-globinopathies, SCD and TM. From the data presented here, one could envision a safe and effective binary therapeutic compound that both induces fetal hemoglobin (by blocking LSD1 activity) and at the same time promotes normal erythroid differentiation by blocking the induction of PU.1 and/or RUNX1. The treatment could also be applicable to the treatment of AML and/or MDS patients suffering from malignant or pre-malignant neoplasms.

## Author Contributions

Conception or design of the work: L.Y., G.M., R.K., K.-C.L., J.D.E.; acquisition and analysis: L.Y., G.M., C-J.K., E.S., Y.W., S.A.S., N.J., K.-C.L.; interpretation of data: L.Y., G.M., C.-J.K., A.W., T.M., R.K., M.Y.; M.G.R., J.P., J.H.B., K.-C.L., J.D.E.; preparation of the draft or substantively revised it: L.Y., G.M., K.-C.L., J.D.E.

## Materials and Methods

### Materials & Correspondence

contact J.D.E. for material requests.

### Statistical information

data are presented as means ± SD by unpaired student’s *t*-test. n.s. = not statistically significant

### Erythroid differentiation of human CD34+ cells

Human CD34+ HSPCs from healthy donors were purchased from Fred Hutchinson Cancer Research Center. The CD34+ cells were cultured and differentiated into mature erythroid cells using a standard three-phase culture system(Giarratana et al., 2011). The basic media was composed of Iscove’s Modified Dulbecco’s Medium (IMDM) supplemented with 5% pooled human AB plasma (Octaplas, Rhode Island Blood Center), 1x penicillin-streptomycin (Gibco), 1% L-glutamine, 330 μg/ml holo-transferrin (T4132, Sigma), 2 IU/ml heparin (H3149, Sigma), and 10 μg/ml human insulin (91077C, Sigma).

From day (d) 0-7 of the expansion culture period, the basic medium was supplemented with 1 μM hydrocortisone (07904, Stem Cell Technologies), 5 ng/ml human IL-3 (203-IL, R&D), 100 ng/ml human SCF (255-SC-050, R&D), and 3 IU/ml erythropoietin (Epoetin Alfa, Amgen). Hydrocortisone and IL-3 were withdrawn from the media beginning on d8 while SCF was omitted after d12. On differentiation day (d) 7, CD34+ HSPC-derived erythroid cells were counted and reseeded at 10^5^ cells/ml in the presence of increasing concentrations of LSD1i (0, 120 nM, 370 nM or 1.1 □M) for an additional 11 days.

The novel LSD1 inhibitor described in this manuscript (CCG50; LSD1i) was designed, synthesized and characterized by the Center for Chemical Genomics at the University of Michigan, and will be described in a separate study (in preparation). At d7 of erythroid differentiation, cells (10^5^/ml) were cultured in the presence of DMSO or 3-fold dilutions of LSD1i (1.1 μM, 370 nM or 120 nM; Fig. S1B). Thereafter, the medium was changed at d11 and d14 with freshly added LSD1i. At the indicated time points, aliquots of cells were removed for flow cytometric, RNA and/or HPLC analyses. Experiments were repeated using CD34+ cells from at least two separate donors.

### *In vitro* LSD1 inhibition assay

A fluorescent *in vitro* LSD1 inhibition assay (Item No. 700120, Cayman Chemicals) was performed according to the manufacturer’s instructions.

### HPLC

On the day erythroid differentiation terminated, cells were washed with PBS and the cell pellets were lysed; the contents were then subjected to HPLC analyses using a BioRad CDM System CDM5.1 VII Instrument according to the manufacturer’s instructions.

### Total RNA isolation and qRT-PCR analyses

Total RNA was isolated from cells using Trizol (ThermoFisher Scientific) or RNeasy Micro Kit (Qiagen) according to manufacturer’s instructions. cDNA was synthesized using SuperScript III Reverse Transcriptase (ThermoFisher Scientific). qRT-PCR was performed using FastSybr Green Mastermix on an ABI Step One Plus instrument (ThermoFisher Scientific). Sequences of all primers are listed in Supplementary Table 1.

### BAC recombination

BAC recombination and the 196 kbp mouse *Gata1* BAC clone (RP23-443E19) have been described previously(Brandt et al., 2008; Takai et al., 2013). Briefly, the targeting DNA fragment was constructed by inserting a CreERT2-polyA-Neo cassette at the translational start site of mouse *Gata1* (in the 2nd exon) between 5’ and 3’ homologous recombination arms. The resultant targeting DNA fragment was excised from the cloning vector and transfected into transiently heat-activated EL250 bacteria (to induce recombinase expression) containing BAC RP23-443E19. Recombinant bacterial colonies (with newly acquired Kanamycin resistance) were individually screened for homologous recombination at the *Gata1* ATG by PCR using two primer sets (Supplemental Table 1). Clones that were determined to have successfully recombined (CreER^T2^ Neo+) were then cultured in the presence of L-arabinose to induce flp recombinase expression to promote excision of the frt-flanked Neo selection marker. Finally, the BAC DNA was purified by NucleoBond BAC100 and submitted for microinjection into fertilized egg of C57BL6 background mice (Transgenic Animal Model core, University of Michigan). Primers are listed in Supplementary Table 1.

### Mice

*G1B*CreER^T2^ transgenic founders were intercrossed with wild type C57BL6 mice for three generations to eliminate possible mosacism and to confirm germline transmission. Two lines, L245 and L259, were interbred with Cre reporter mice R26 tdTomato (Madisen et al., 2010) (Jackson Stock# 007914) or homozygous *Lsd1*^f/f^ mutant mice(Wang et al., 2007). To induce *G1B*CreER^T2^ activity, mice (from 8-12 weeks old) were injected intraperitoneally with tamoxifen (Tx; 2 mg per injection on alternate days for 2 weeks). All animal procedures were approved by the Institutional Animal Care and Use Committee of the University of Michigan (IUCAC Protocol PRO00009778). Primer sequence for genotyping are listed in Supplemental Table 1.

### Differential peripheral blood analyses

Peripheral blood was collected from mice by facial vein bleed into EDTA-treated tubes and analyzed on a Hemavet according to the manufacturer’s instructions.

### Genomic DNA qPCR

Genomic DNAs were purified from FACS-sorted mouse basophilic erythroblasts. To quantify *Lsd1* deletion efficiency, qPCR was performed using FastSybr Green Mastermix (Applied Biosystems) and *Lsd1* primer pairs that specifically annealed to genomic sequences within (P1) or outside of (P2) the two LoxP sites flanking *Lsd1* exon 6 (Wang et al., 2007)(Fig. 3H). Deletion efficiency was calculated from the Ct ratio of P1/P2 as described previously(Wang et al., 2007; Yu et al., 2014). The primers used for qPCR analysis are listed in Supplemental Table 1.

### Flow cytometry

Total bone marrow (BM) isolation and subsequent antibody staining protocols were described previously(Moriguchi et al., 2015). AnnexinV staining was performed as described previously (Yu et al., 2017). Flow analysis was performed on a BD Fortessa while flow sorting was performed on a BD Aria III. Antibodies used for flow cytometry are listed in Supplemental Table 2.

### Chromatin Immunoprecipitation

ChIP assays were conducted as described previously(Yu et al., 2018).

### Colony-Forming Unit (CFU) assay

Isolated total BM cells or purified, sorted cell populations were counted and seeded into Methocult M3434 or M3334 (Stemcell Technologies) according to manufacturer’s instructions for enumeration of BFU-E, CFU-GM, CFU-GEMM or CFU-E colonies.

### CRISPR knockout of *Pu.1* or *Runx1* genes

Genome edited *Pu.1* or *Runx1* loss of function strategy was essentially as described in a previous study(Yu et al., 2018). Individual HUDEP2 *Pu.1* or *Runx1* targeted clones were initially identified by PCR (sequences shown in Supplemental Table 1) and verified by Sanger sequencing. For each gene, four clones were generated using two different sgRNAs. Properly targeted mutation of all clones (except one) was confirmed by Sanger sequencing (Supplemental Figs. 12, 13). Three different nontargeting sgRNAs (NT1, NT4 and NT3) containing scrambled guide sequences were used to generate negative control HUDEP2 clones. One clone (P2-4) exhibited no amplification of the guide-targeted exon, suggesting that this clone bore a large genomic DNA deletion (data not shown). The guide sequences used for gene editing are also listed in supplemental Table 1.

## Acknowledgements

We gratefully acknowledge the assistance and insights provided by multiple colleagues at the University of Michigan (Sojin An, Uhnsoo Cho, Susan Hagen, Pil Li and Mathivanan Packiarajan). We gratefully acknowledge continued support from the NHLBI (awards U01 HL117658 and P01 HL146372 to J.D.E. and A.W.), from the Cooley’s Anemia Foundation (L.Y.) as well as a center of excellence award from the NIDDK (U54 DK106829) to the Fred Hutchinson Cancer Center to support the isolation and distribution of normal human hematopoietic stem and progenitor (CD34+) cells for the scientific community.

## Supplemental Information

**Supplemental Figure 1:**
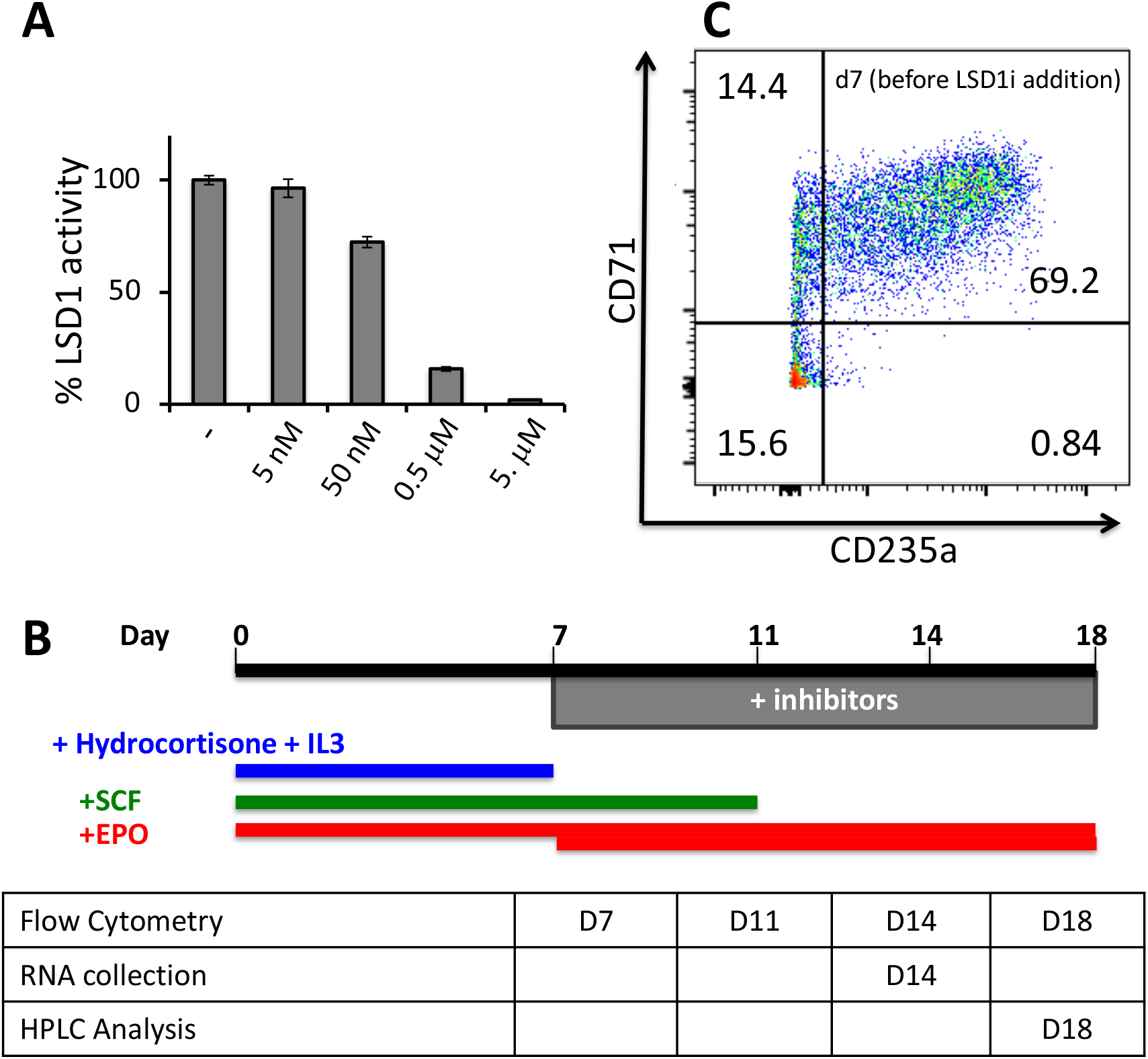
Characterization of Inhibitor Effects on Human CD34+ HSPC-induced Erythroid Differentiation. (A) LSD1 inhibition by a novel, reversible LSD1 inhibitor, CCG50 (LSDi). (B) Schematic summarizing the timing and duration of additives (hydrocortisone, IL3, SCF and EPO) media supplements during *in vitro* erythroid differentiation of human CD34+ HSPCs. Hydrocortisone and human IL-3 were withdrawn on day7 when the LSD1 inhibitor(s) were added; human SCF was withdrawn from culture media on day11. (C) Representative flow cytometric cell count of CD34+ HSPCs undergoing erythroid differentiation at d7 (prior to LSD1 inhibitor treatment) analyzed with anti-CD71 and anti-CD235a erythroid antibodies.

**Supplemental Figure 2:**
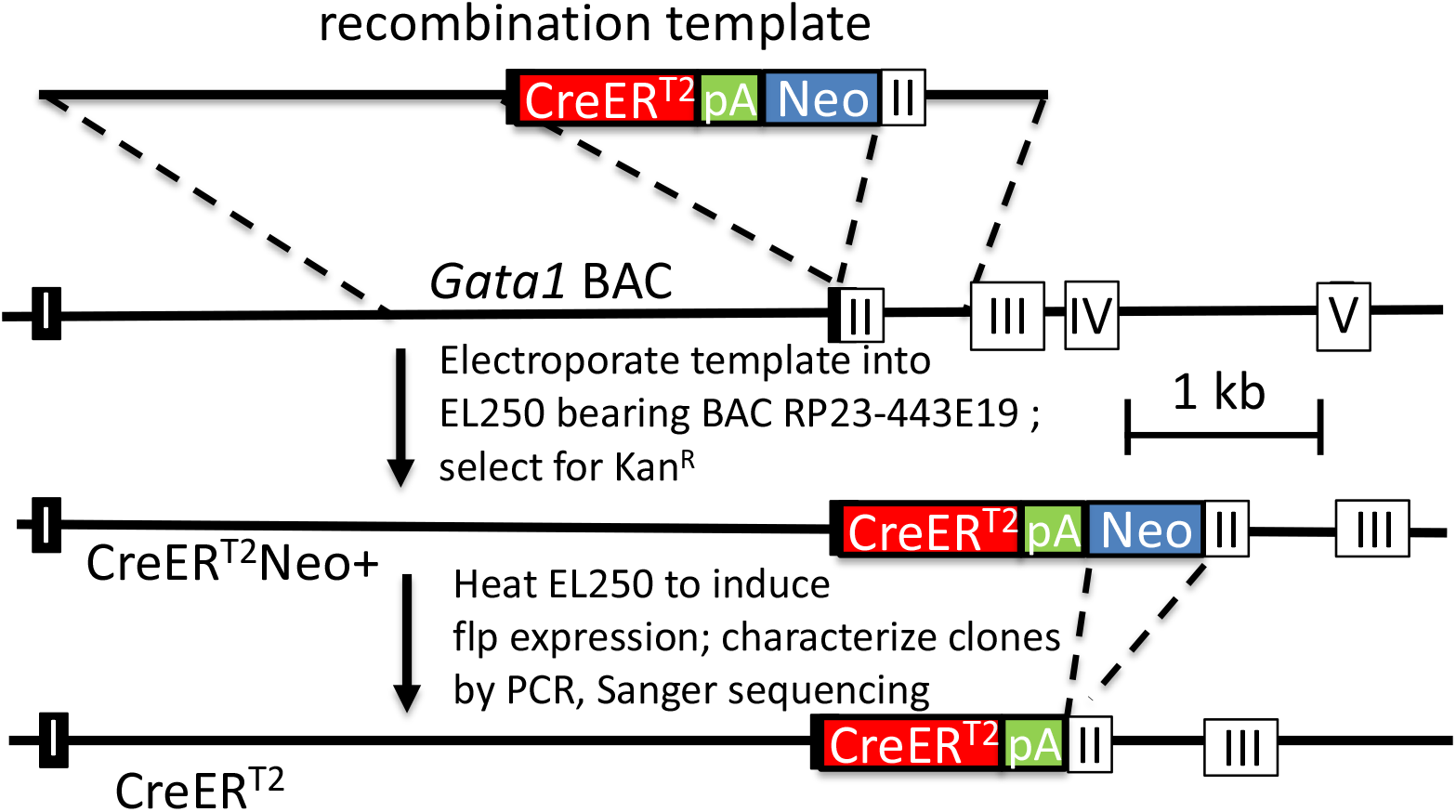
Mouse *Gata1* BAC recombination strategy. A creER^T2^-PolyA-Neo cassette (Takai et al. 2013; Yu et al. 2017) was targeted to the ATG start codon in exon 2 of a 196 kb wild type mouse *Gata1* BAC (*G1B*). Subsequently, the Neo selection cassette was removed from the recombinant *G1B*CreER^T2^-Neo BAC to generate the resultant *G1B*CreER^T2^ BAC, which was used to generate two lines of transgenic mice.

**Supplemental Figure 3:**
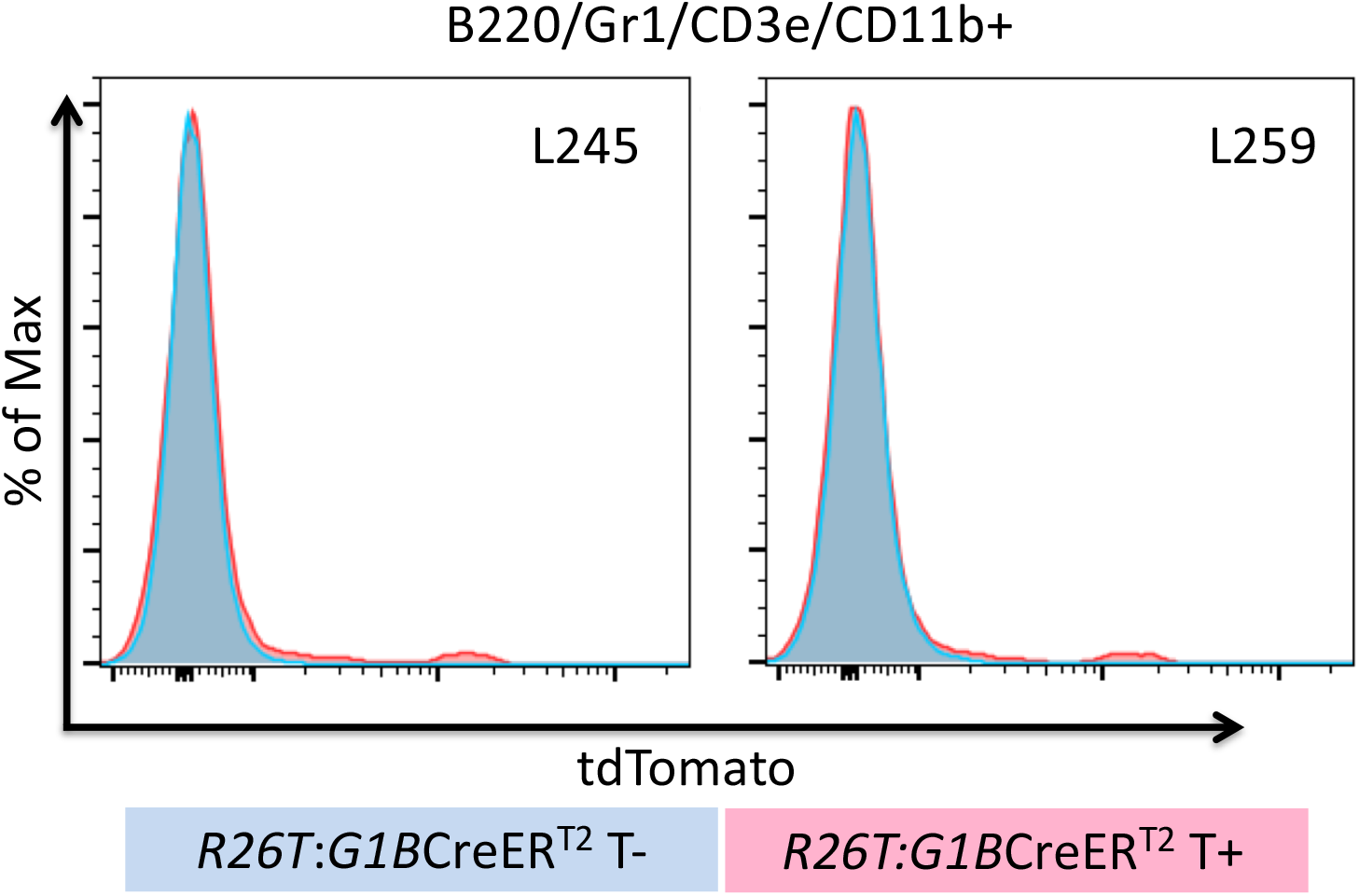
*G1B*CreERT2 is not active in non-erythroid lineages. TdTomato epifluorescence was not detectable in B220+, Gr1+, CD11b+ or CD3e+ BM cells of *G1B*CreER^T2^ lines L245 and L259 treated with Tamoxifen (red peak). R26T:*G1B*CreER^T2^ mice that were not treated with Tx served as negative controls (blue peak).

**Supplemental Figure 4:**
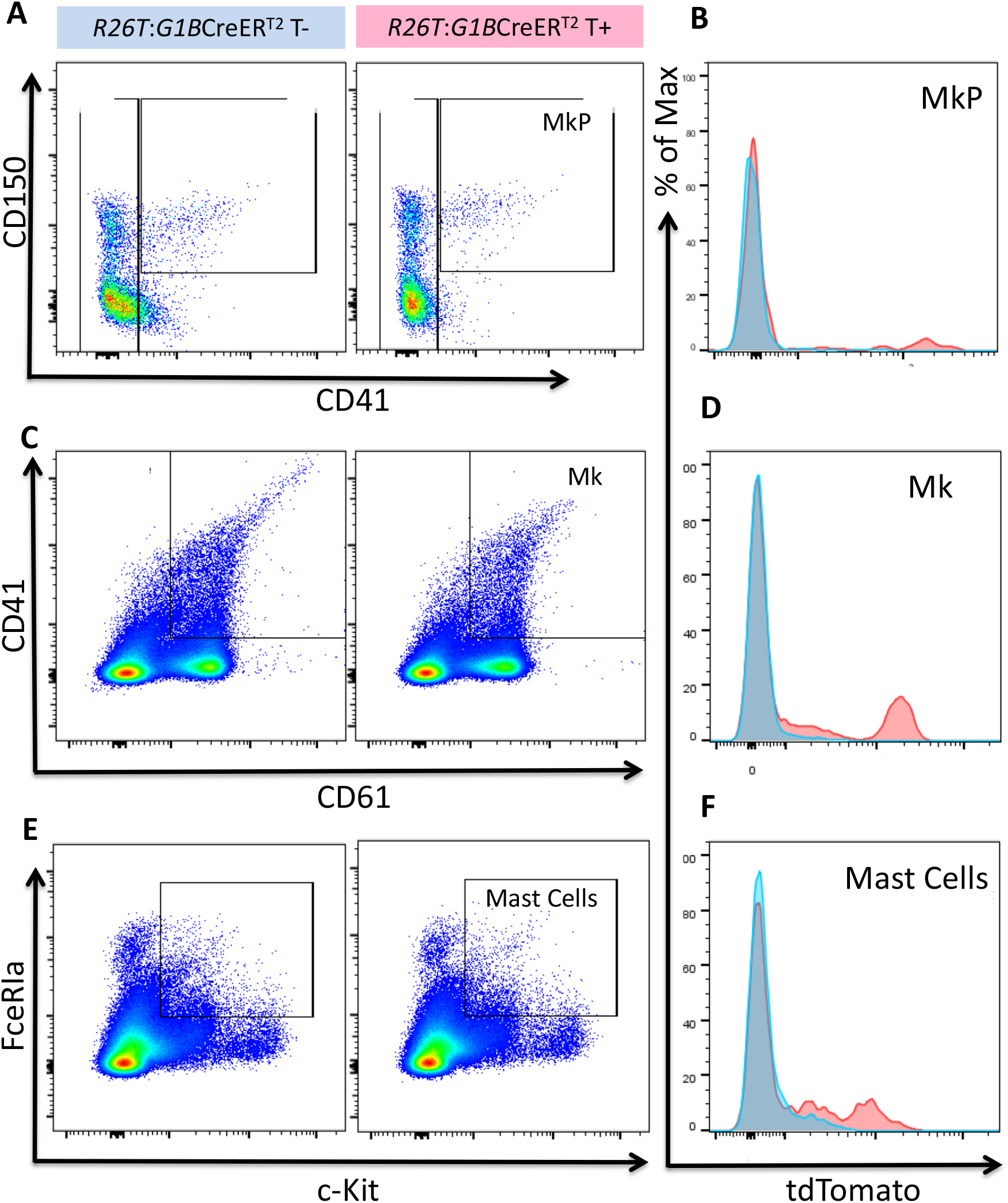
*G1B*CreER^T2^ is expressed in a minor fraction of megakaryocytes and mast cells. Representative flow cytometric plots depicting the gating cutoffs for megakaryocytic progenitor cells (MkPs; Lin-cKit+Sca1-CD41+CD150+; A, B), megakaryocytes (Mk; CD61+CD41+; C, D) and mast cells (FceRIa+c-Kit+; E, F) in the BM of untreated (left panels in A, C and E; blue peaks in B, D and F) or Tx-treated (right panels in A, C and E; red peaks in B, D and F) R26T:*G1B*CreER^T2^ L259 mice. Representative flow histograms depict TdTomato (red peaks) in very few MkPs (B), Mk (D) or mast cells (F).

**Supplemental Figure 5:**
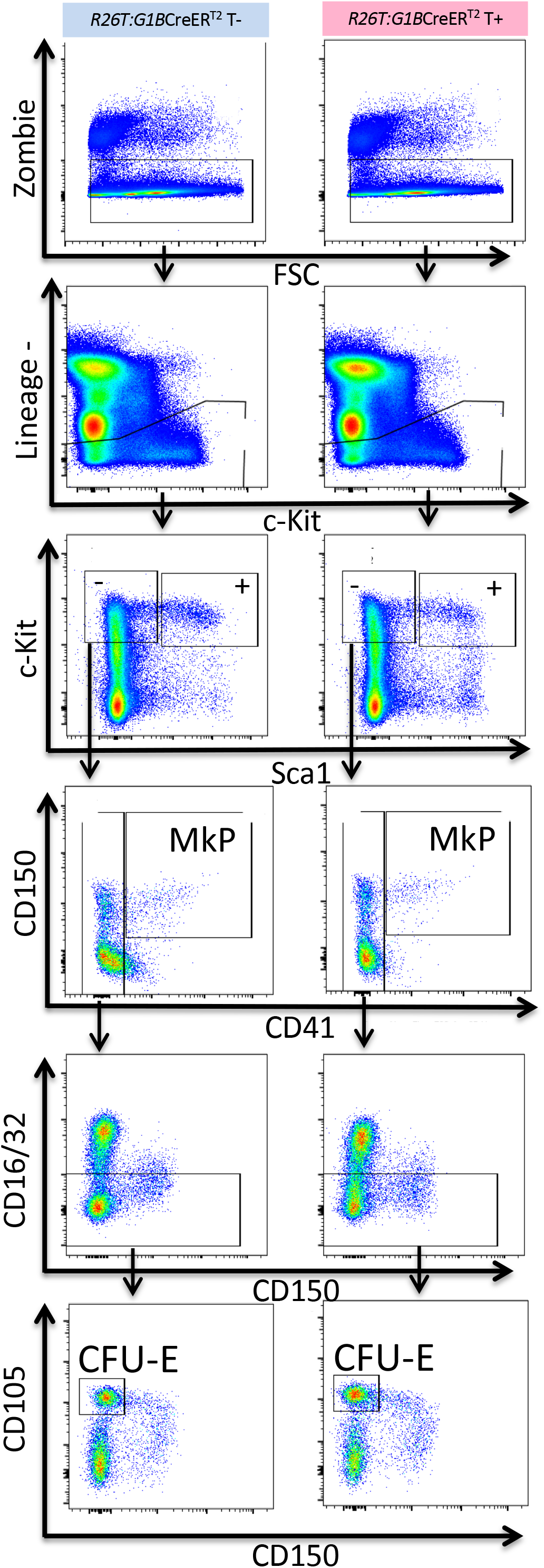
Gating strategies for CFU-E and megakaryocytic progenitor cell populations. Total BM cells from untreated (left panels) or Tx-treated (right panels) R26T:*G1B*CreER^T2^ L259 mice were stained with the indicated lineage antibodies. Representative flow plots showing gating cutoffs for CFU-Es (Lin-cKit+Sca1-CD41-CD16/32- CD150-CD105+) and megakaryocytic progenitors (MkPs; Lin-cKit+Sca1-CD41+CD150+), which were analyzed for TdTomato epifluorescence in Fig.s 2C and S4, respectively.

**Supplemental Figure 6:**
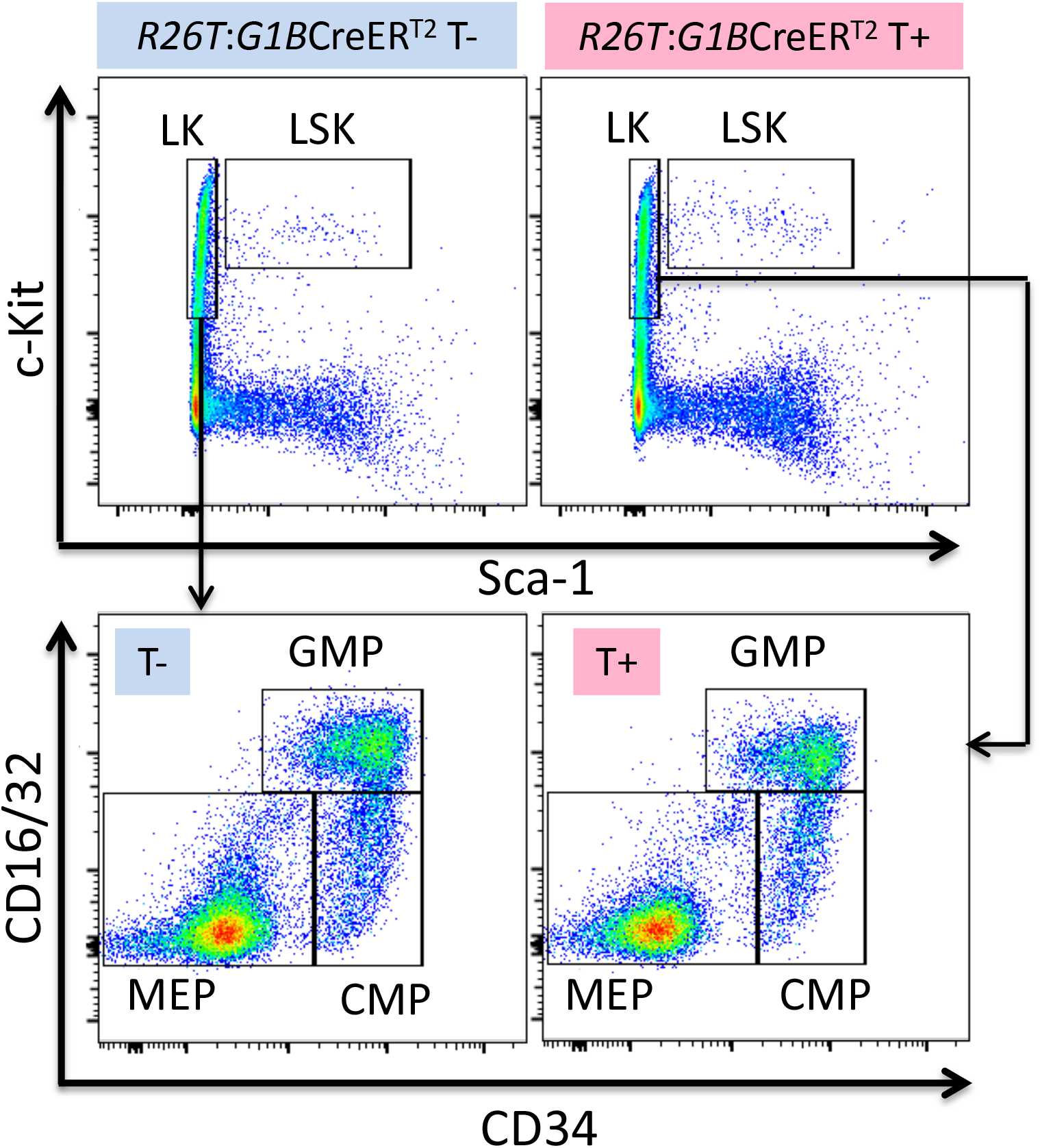
Gating strategies for LSK, CMP, GMP and MEP cell populations. Total BM cells from untreated (left panels) or Tx-treated (right panels) R26T:*G1B*CreER^T2^ L259 mice were stained with Lineage-, c-Kit-, Sca1-, CD34- or CD16/32 antibodies. Representative flow plots showing gating strategies defining the LSK (Lin-c-Kit+Sca1+), CMP (Lin-c-Kit+Sca1-CD16/32-CD34+), GMP (Lin-c-Kit+Sca1-CD16/32+CD34+) and MEP (Lin-c-Kit+Sca1-CD16/32-CD34-) populations, which were further analyzed for TdTomato epifluorescence in Fig. 2C.

**Supplemental Figure 7:**
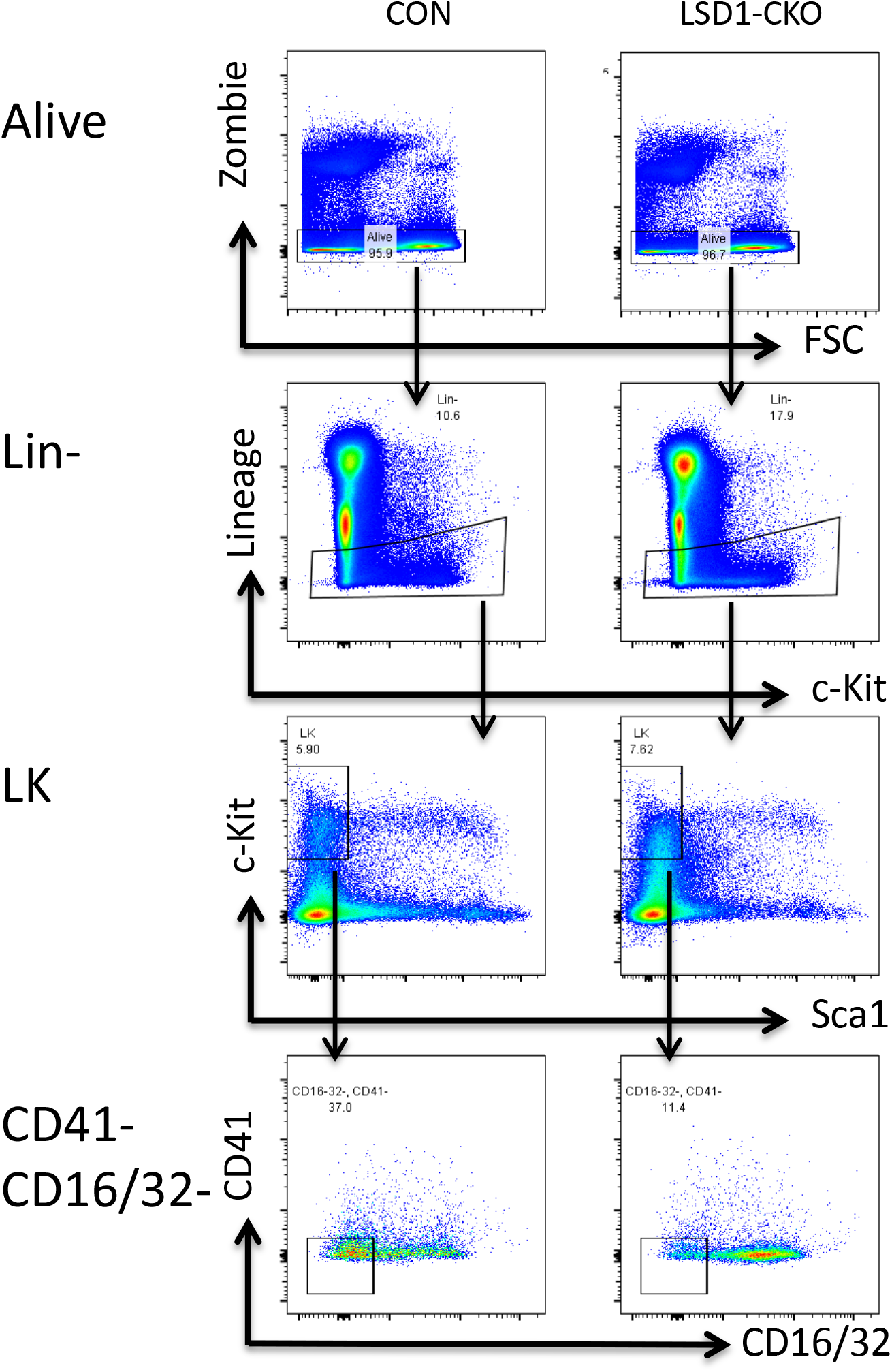
Gating strategies for CMP, GMP and MEP populations. The CD41-CD16/32- LK fraction was separated by CD105 and CD150 for CFU-E staining as shown in Figure 3. The LK fraction was separated by CD34 and CD16/32 for CMP, GMP and MEP staining in Figure 4.

**Supplemental Figure 8:**
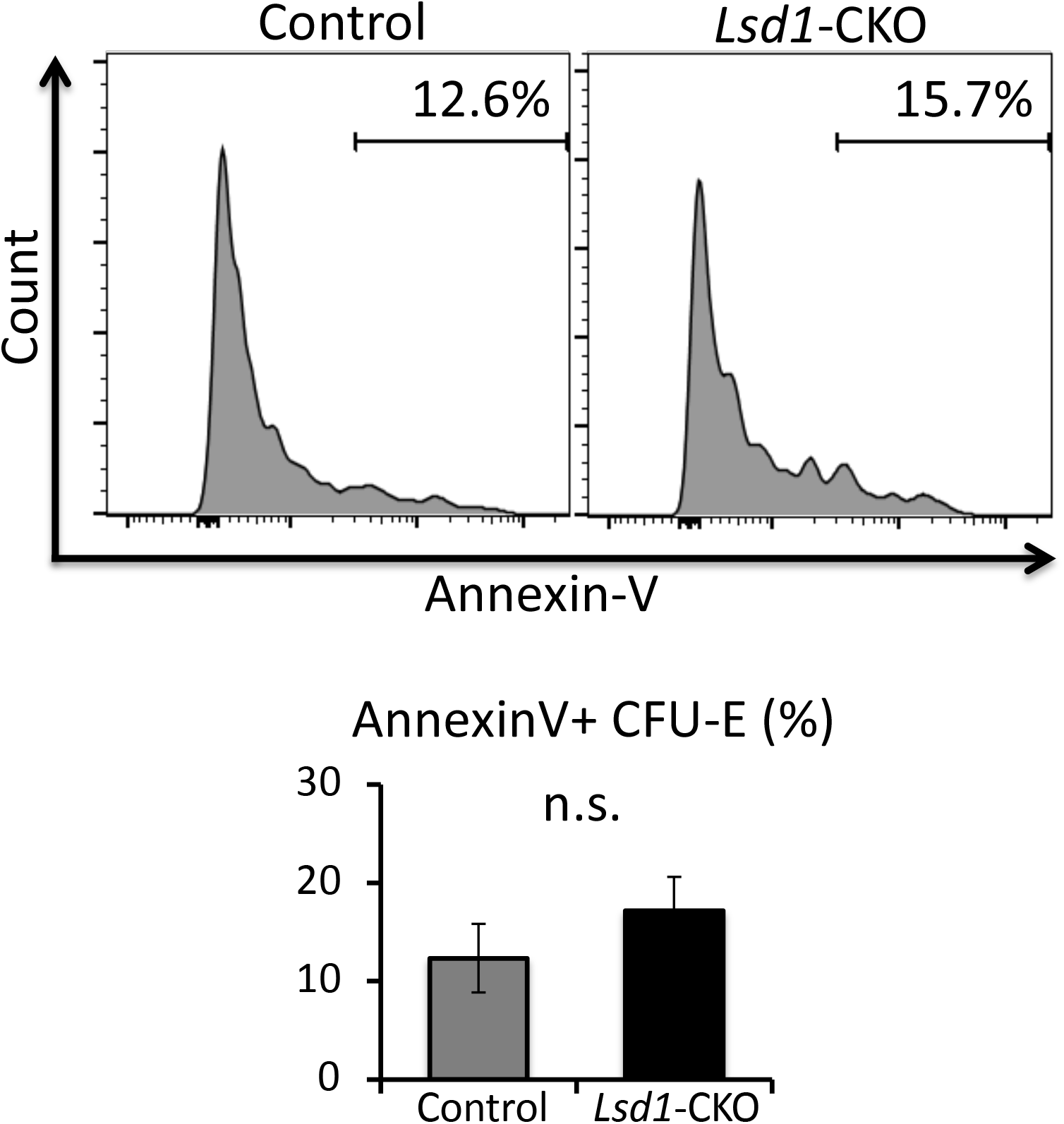
CFU-E cell death is comparable in *Lsd1-CKO* and control mice. The cell death of CFU-E cells in Figure 3D was analyzed by AnnexinV staining.

**Supplemental Figure 9:**
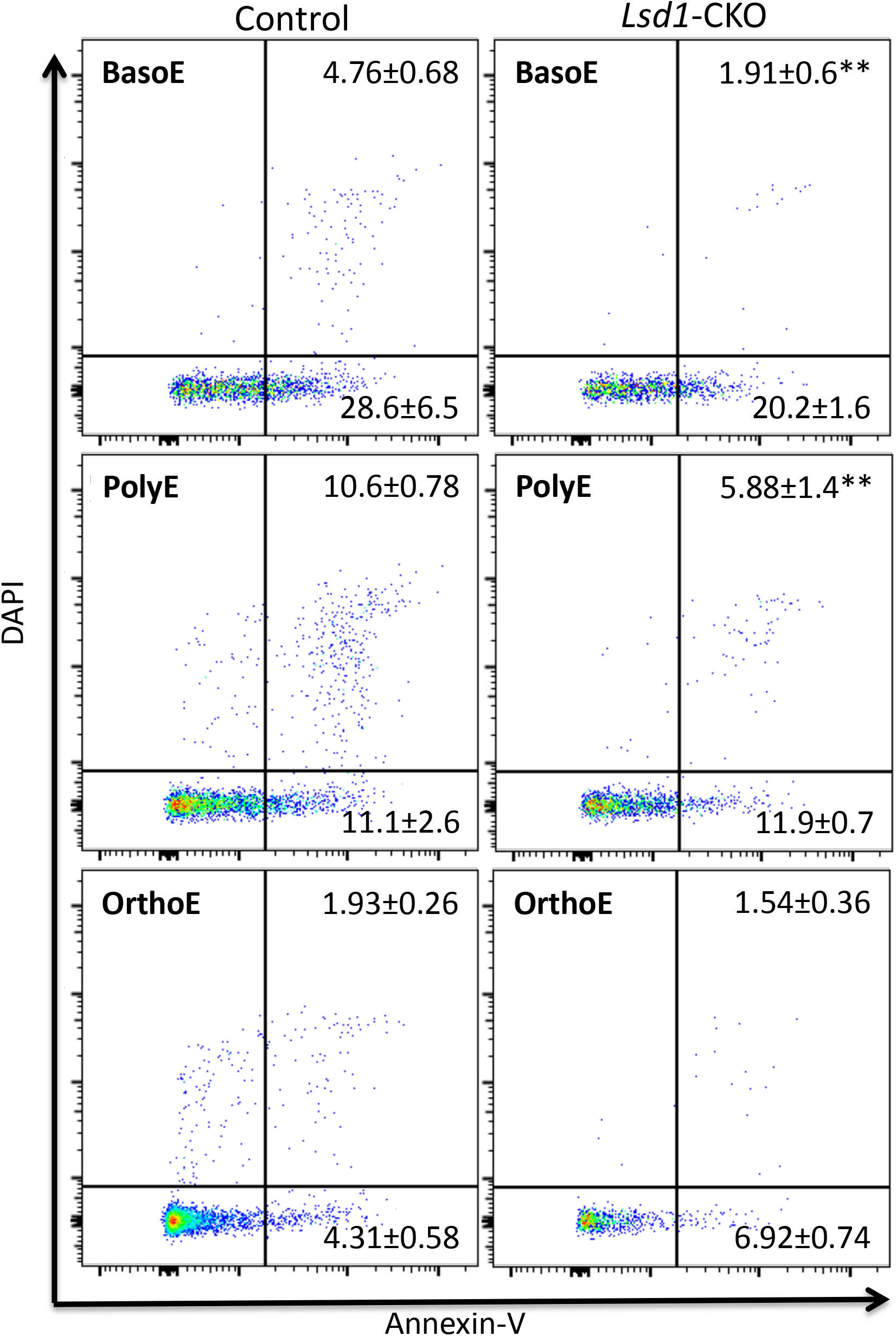
Erythroid precursor cells in *Lsd1*-CKO mice have less cell death. The apoptotic cell death of BasoE, PolyE and OrthoE in Fig. 3F was assessed by AnnexinV staining. Data are shown as the means ± SD. (** *p*<0.01; unpaired Student’s t-test).

**Supplemental Figure 10:**
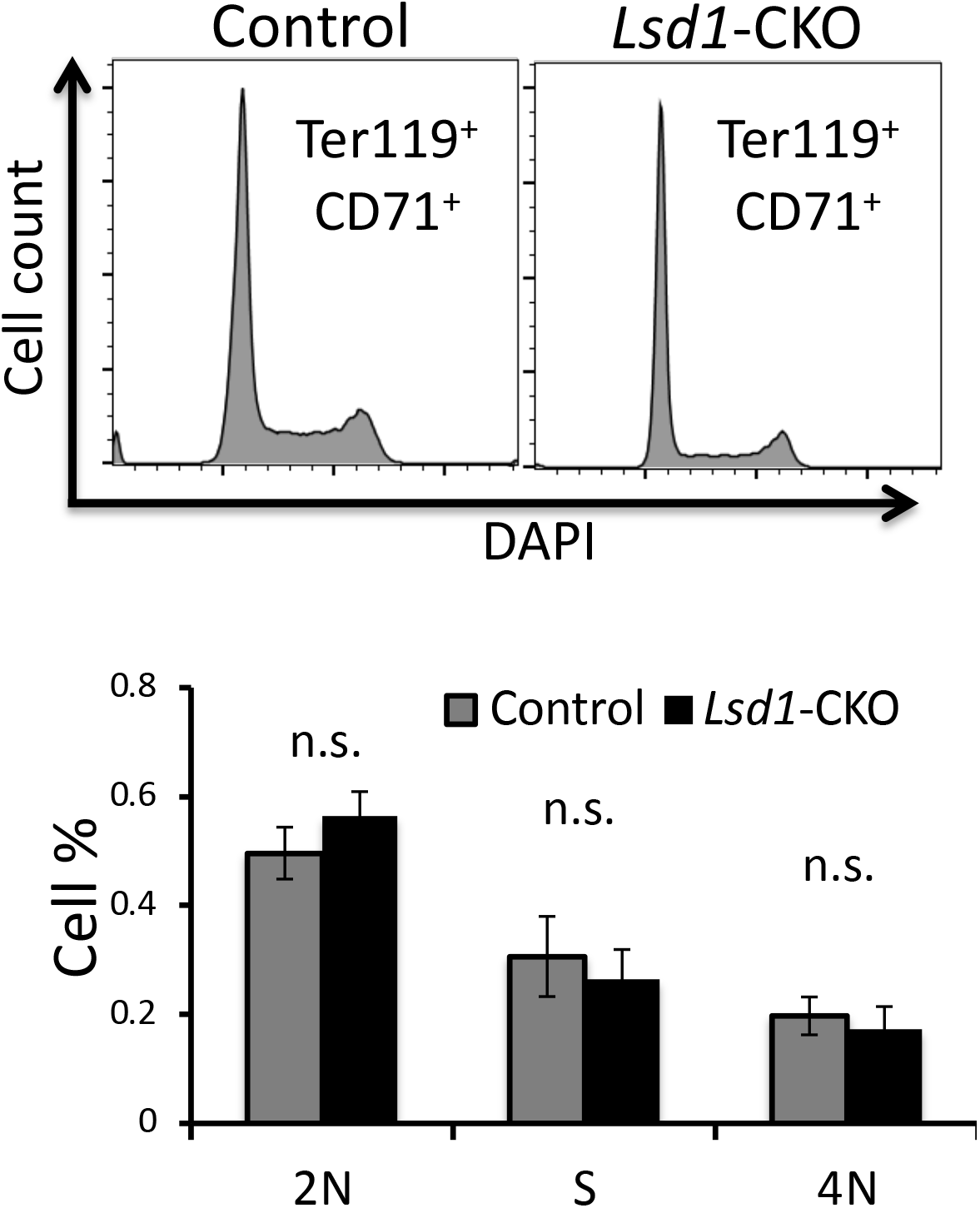
The cell cycle status of erythroid precursors in *Lsd1*-CKO and control mice is comparable. The distribution of CD71+Ter119+ cells depicted in Figure3F was analyzed by DAPI staining. Data is shown as the means ± SD. (** *p*<0.01; unpaired Student’s t-test). 2N, cells in G1 phase of cell cycle; S, cells in S phase of cell cycle; 4N, cells in G2/M phase of the cell cycle.

**Supplemental Figure 11:**
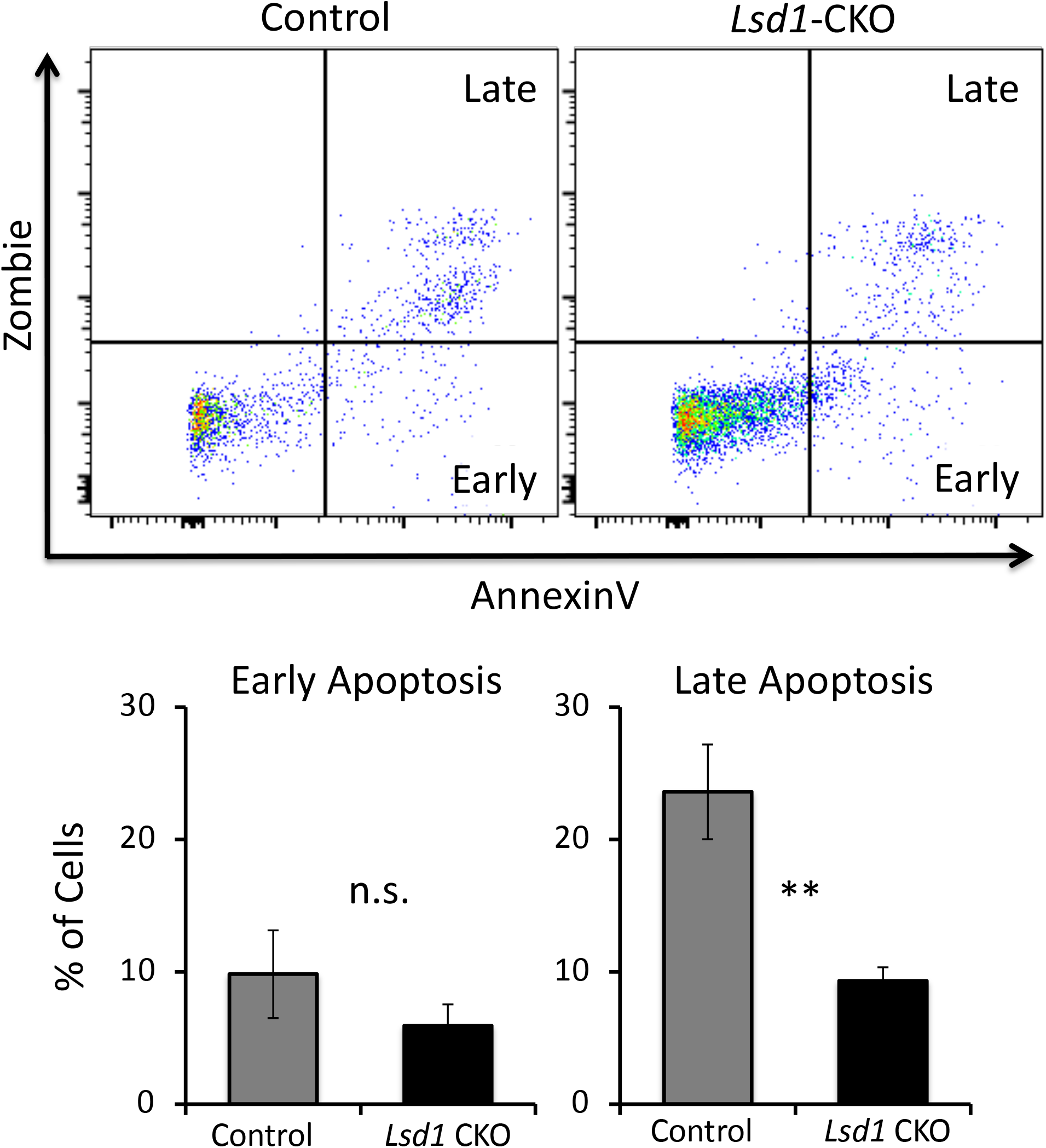
GMP in *Lsd1*-CKO mice have significantly diminished cell death. The cell death of GMP shown in Figure 4A was analyzed by AnnexinV staining. Data was shown as the means ± SD. (** *p*<0.01; unpaired Student’s t-test).

**Supplemental Figure 12:**
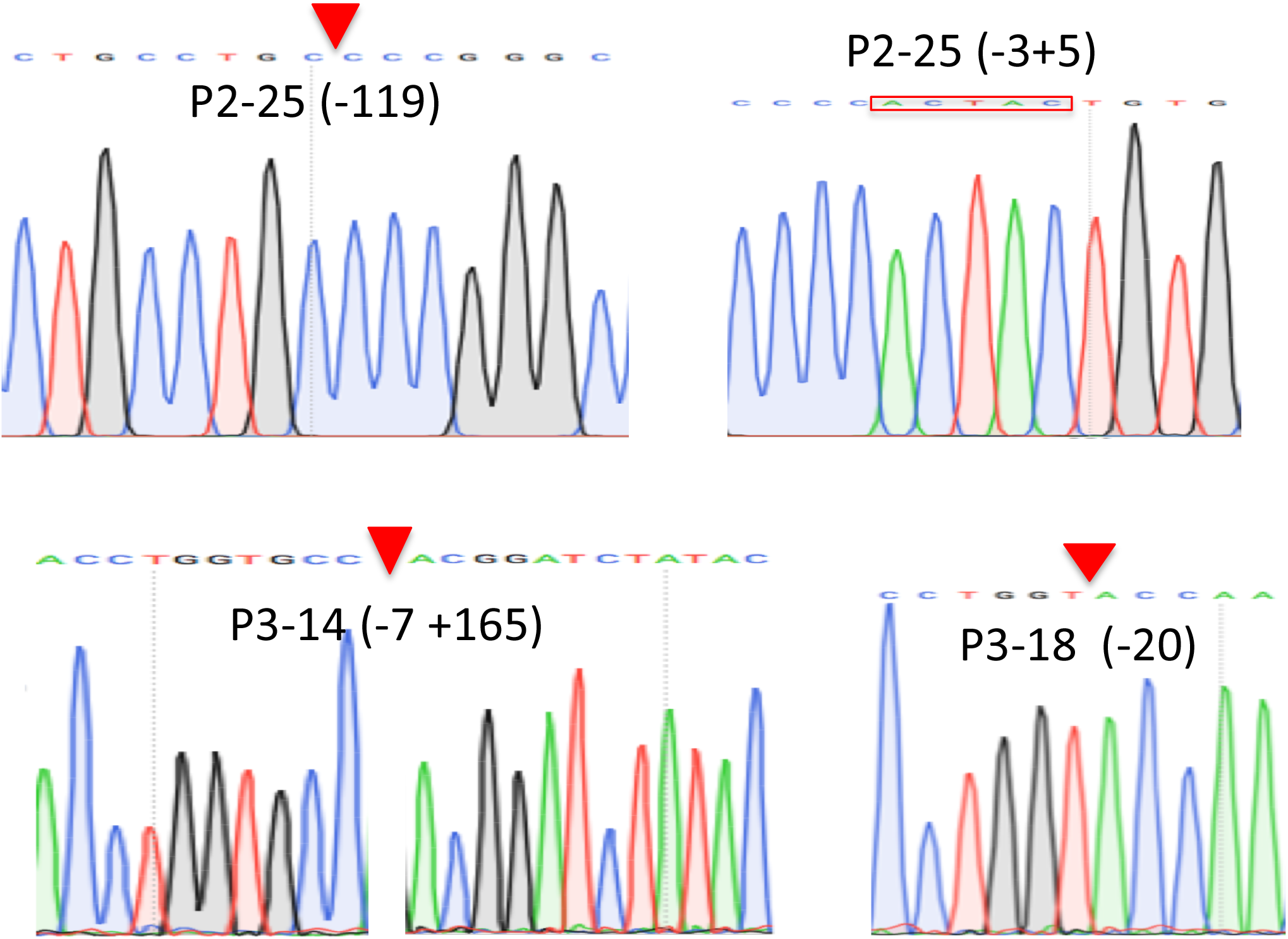
Sanger sequencing data of *Pu.1*-/- HUDEP2 clones. Individual HUDEP2 clones were subjected to genomic DNA extraction, the genomic DNA locus of sgRNA cutting sites were amplified by PCR, the amplicons were subcloned into sequencing vector and the sanger chromatogram data surrounding the cutting sites were shown. Note, clone P2-25 has two PCR products with different size; clone P3-14 and P3-18 have only one PCR product; and clone P2-4 has no PCR product, all PCR amplicons were sequenced.

**Supplemental Figure 13:**
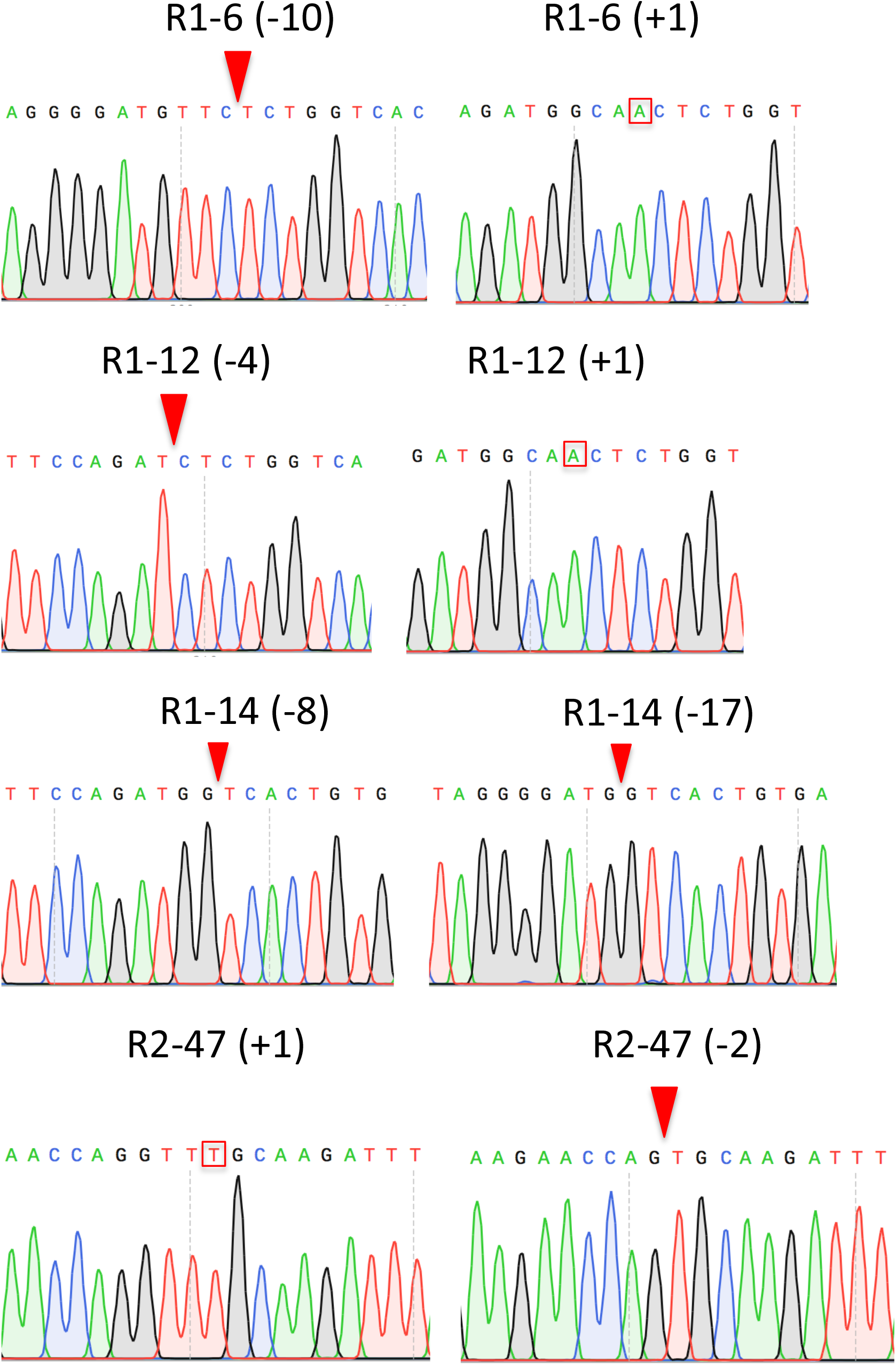
Sanger sequencing data of Runx1-/- clones. Each individual clones were subjected to genomic DNA extraction, the genomic DNA locus of sgRNA cutting sites were amplified by PCR, the amplicons were subcloned into sequencing vector and the sanger chromatogram data surrounding the cutting sites were shown.

**Supplemental Figure 14:**
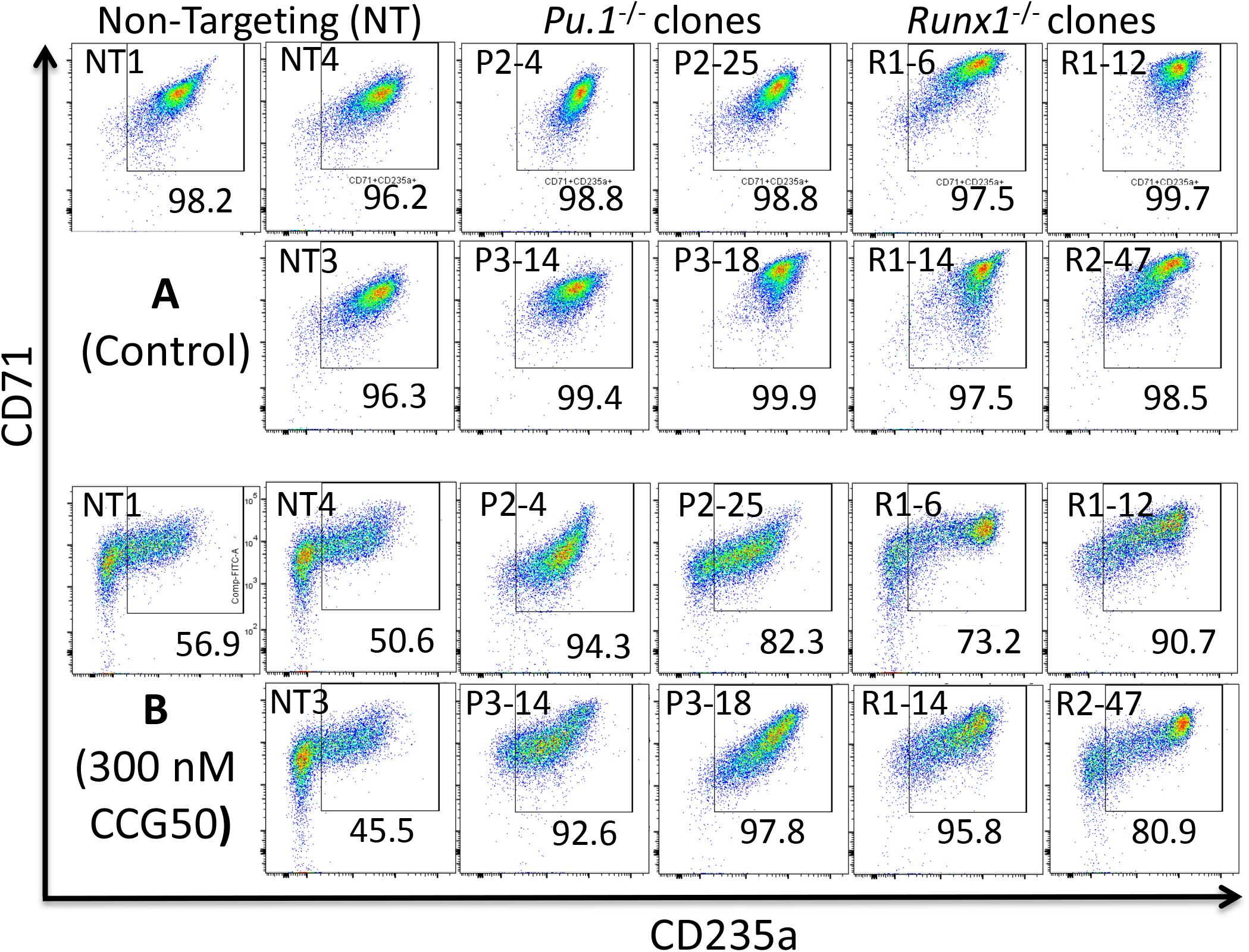
*Pu.1* or *Runx1* knockout rescues HUDEP2 cells from the side effects of LSD1 inhibition. CD71/CD235a flow cytometry analysis of three different nonspecific sgRNA-infected cells (left), four *Pu.1-/-* clones (middle) or four *Runx1-/-* clones (right) treated with **(A)** DMSO or **(B)** 300 nM CCG50 LSD1i for four days after induction of erythroid differentiation.

**Supplemental Figure 15.**
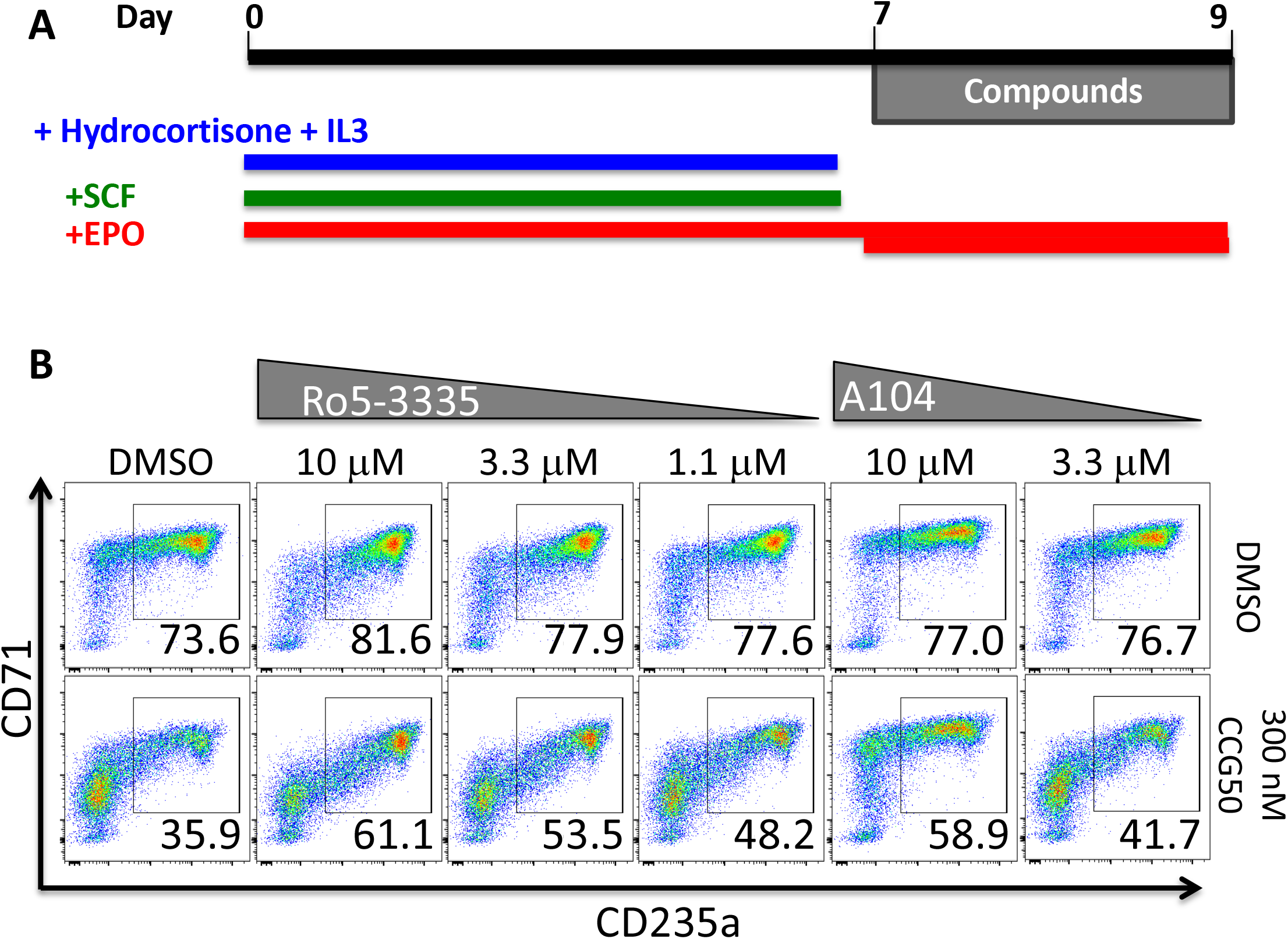
RUNX1 inhibitor co-treatment partially rescues the toxicity of LSD1 inhibitor CCG50. **(A)** Human CD34+ HSPC were expanded for 7 days; at day7, cells were reseeded and treated with LSD1 inhibitor CCG50, RUNX1 inhibitors (Ro5 or A-10-104) or inhibitors of both factors. **(B)** CD71/CD235a staining of differentiating human CD34+ cells in erythroid differentiation at day 9, 2 days after adding CCG50, RUNX1 inhibitor Ro5-3335, RUNX1 inhibitor AI-10-104 (A104) or both LSD1 and RUNX1 inhibitors.

**Supplemental Table 1.**
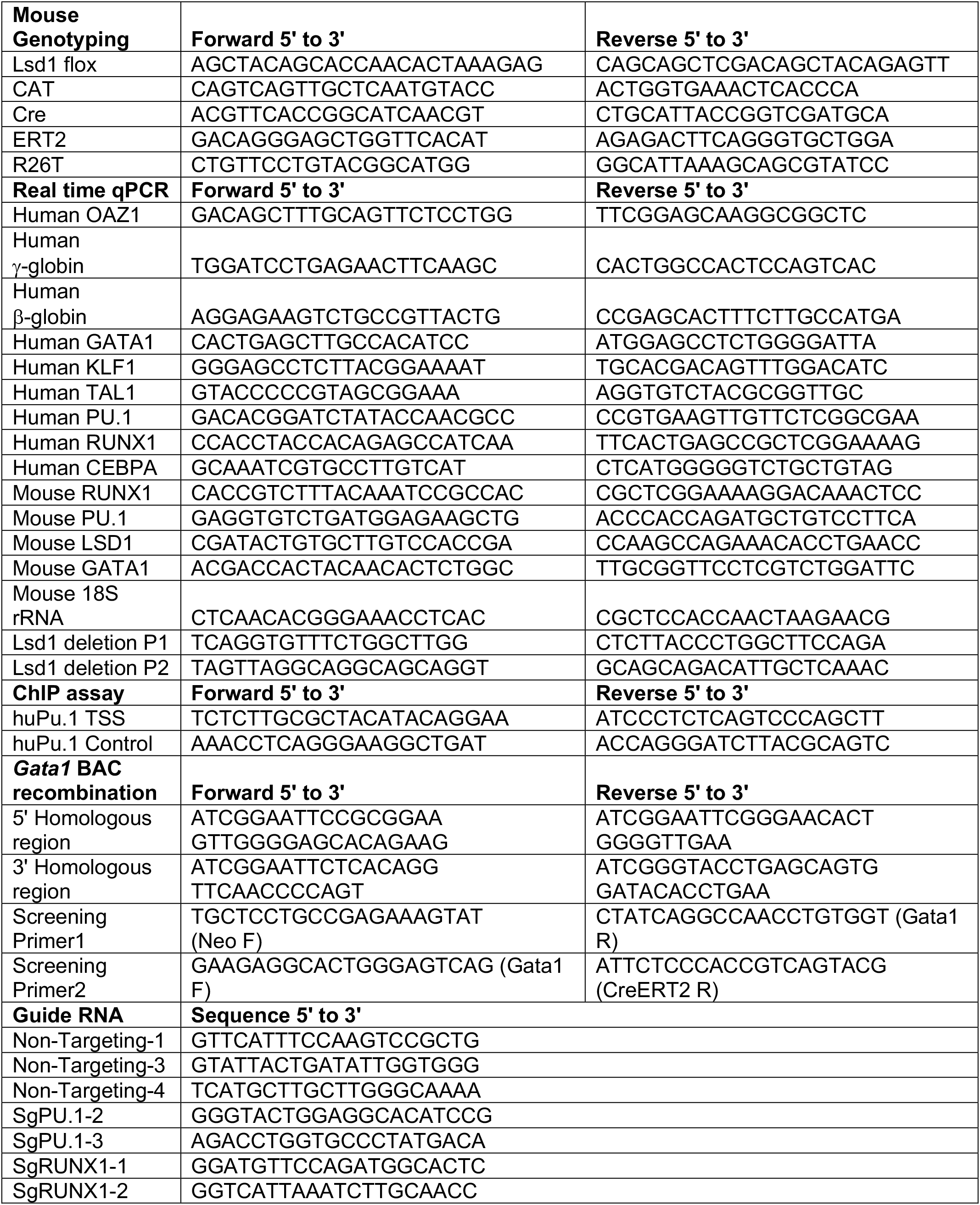
Sequences of all DNA oligonucleotides used in these studies,

**Supplemental Table 2.** List of all antibodies, antigens recognized, and the suppliers used in these studies,

| **Human** | **Fluorescence** | **Cat#** | **Vendor** |
| --- | --- | --- | --- |
| CD235a | APC | 551336 | BD Biosciences |
| CD71 | FITC | 334104 | BioLegend |
| CD11b | PE | 301305 | BioLegend |
| **Mouse** | **Fluorescence** | **Cat#** | **Vendor** |
| CD71 | FITC | 113806 | BioLegend |
| Ter119 | APC | 116212 | BioLegend |
| CD44 | APC-Cy7 | 103028 | BioLegend |
| B220 | FITC | 103206 | BioLegend |
| Gr1 | FITC | 108406 | BioLegend |
| CD3e | FITC | 100306 | BioLegend |
| CD11b | FITC | 101205 | BioLegend |
| CD150 | APC | 115910 | BioLegend |
| CD41 | Alexa Fluor 700 | 133925 | BioLegend |
| CD41 | FITC | 133904 | BioLegend |
| CD61 | APC | 104315 | BioLegend |
| FceRIa | FITC | 134305 | BioLegend |
| c-Kit | APC | 105812 | BioLegend |
| B220 | Biotin | 13-0452-85 | ThermoFisher |
| CD2 | Biotin | 100103 | BioLegend |
| CD3 | Biotin | 100304 | BioLegend |
| CD5 | Biotin | 100604 | BioLegend |
| CD8a | Biotin | 100704 | BioLegend |
| GR1 | Biotin | 13-5931-85 | ThermoFisher |
| TER119 | Biotin | 116204 | BioLegend |
| StAV | BV650 | 405232 | BioLegend |
| c-Kit | APC-Cy7 | 105825 | BioLegend |
| Sca1 | BV421 | 108128 | BioLegend |
| CD16/32 | BV605 | 563006 | BD Biosciences |
| CD105 | PE-Cy7 | 120410 | BioLegend |
| Sca1 | APC | 108111 | BioLegend |
| CD16/32 | PE-Cy7 | 101317 | BioLegend |
| CD34 | FITC | 553733 | BD Biosciences |
| Ter119 | eF450 | 48-5921-82 | ThermoFisher |
| Gr1 | eF450 | 48-5931-80 | ThermoFisher |
| B220 | eF450 | 48-0452-82 | ThermoFisher |
| CD3e | eF450 | 48-0032-82 | ThermoFisher |
| CD11b | eF450 | 48-0112-80 | ThermoFisher |
| CD41 | eF450 | 48-0411-82 | ThermoFisher |
| **Live/Dead** | **Fluorescence** | **Cat#** | **Vendor** |
| Annexin-V | PE | 556421 | BD Biosciences |
| DAPI | Pacific Blue | 422801 | BioLegend |
| Zombie Aqua | BV510 | 423101 | BioLegend |

